# Modeling soybean growth: A mixed model approach

**DOI:** 10.1101/2023.06.13.544713

**Authors:** Maud Delattre, Yusuke Toda, Jessica Tressou, Hiroyoshi Iwata

## Abstract

The evaluation of plant and animal growth, separately for genetic and environmental effects, is necessary for genetic understanding and genetic improvement of environmental responses of plants and animals. We propose to extend an existing approach that combines nonlinear mixed-effects model (NLMEM) and the stochastic approximation of the Expectation-Maximization algorithm (SAEM) to analyze genetic and environmental effects on plant growth. These tools are widely used in many fields but very rarely in plant biology. During model formulation, a nonlinear function describes the shape of growth, and random effects describe genetic and environmental effects and their variability. Genetic relationships among the varieties were also integrated into the model using a genetic relationship matrix. The SAEM algorithm was chosen as an efficient alternative to MCMC methods, which are more commonly used in the domain. It was implemented to infer the expected growth patterns in the analyzed population and the expected curves for each variety through a maximum-likelihood and a maximum-a-posteriori approaches, respectively. The obtained estimates can be used to predict the growth curves for each variety. We illustrate the strengths of the proposed approach using simulated data and soybean plant growth data obtained from a soybean cultivation experiment conducted at the Arid Land Research Center, Tottori University. In this experiment, plant height was measured daily using drones, and the growth was monitored for approximately 200 soybean cultivars for which whole-genome sequence data were available. The NLMEM approach improved our understanding of the determinants of soybean growth and can be successfully used for the genomic prediction of growth pattern characteristics.

**Author summary:** Nonlinear models are useful for modeling animal and plant growth; however, their parameters are influenced by both genetic and environmental factors. If the same model can be applied to data with different genetic and environmental factors by allowing parameter variations, it can be used to understand, predict, and control the genetic and environmental influences of growth models based on parameter variation. In this study, we propose a statistical method based on nonlinear mixed-effects modeling. The simulation and real data analysis results show that the proposed method was effective in modeling the growth of genetically different soybean varieties under different drought conditions. The usefulness of the proposed method is expected to increase, as high-throughput measurements provide growth data for a large number of genotypes in various environments.

## 1 Introduction

Controlling and predicting the growth patterns of plants and animals are important for growing crops and raising livestock. For example, in crops, growth patterns are known to be closely related to the amount of carbon fixation and thus harvested product [1]. Plant growth patterns are determined by genetics and the environment, and if we could model the relationship between genetics, environment, and growth patterns, we can control and predict the growth based on that model; for example, by properly managing the cultivation of varieties with desirable growth patterns, which can have a positive impact on the yield and quality of the harvest. From this perspective, many recent attempts have been made to measure the growth over time and to model the growth data obtained. In particular, in crop science, the development of remote-sensing technologies, such as unmanned aerial vehicles [2] and proximal sensing [3, 4], has enabled the modeling of the growth curves of many genotypes simultaneously [5–7].

As mentioned earlier, growth patterns are determined not only by genetics and the environment but also by physiological constraints. Owing to physiological constraints, growth patterns often exhibit a nonlinear pattern in which growth is slow in the early stages, accelerates in the mid-term, but slows down after a certain period, and reaches a plateau in the late stages. Such patterns are often approximated by nonlinear functions such as a logistic curve [8] or a Gompertz curve [9]. Because growth patterns are determined by genetic and environmental factors, the parameters of the nonlinear functions describing the growth patterns change under the influence of these factors. Based on this perspective, modeling was conducted in a two-step process: the growth patterns were first approximated by a nonlinear function, and thereafter the estimated parameters of the function were modeled based on the effects of genetic and environmental factors [10–13]. For example, Toda et al. (2022) applied a two-step process to growth data (changes in leaf area index) of various soybean cultivars grown under different growing conditions [13]. They first approximated the observed growth using a nonlinear function and then constructed a model in which the parameters estimated for the nonlinear function were explained by the genetic effects of genome-wide marker genotypes and variations due to blocks. Modeling the genetic effects of genome-wide marker genotypes is a technique known as genomic prediction [14], and these predictions are used to select superior genotypes in plant and animal breeding. Toda et al. (2022) combined the genomic prediction potential of nonlinear functional parameters with early growth data under a Bayesian framework and showed that such a combination could accurately predict growth patterns in later growth stages.

The two-step approach described above is relatively easy to apply but has a drawback. The first step approximates the growth of each individual or plant separately with a nonlinear function; however, this approximation does not consider the factors behind the parameters of the nonlinear model, that is, genetic and environmental effects. For example, given a pair of genetically related samples, these samples are likely to exhibit similar growth and the parameters of the nonlinear model are likely to exhibit similar values. In the second stage of modeling, such factors that are ignored in the first stage are also considered [15]. If the underlying genetic and environmental factors are considered during nonlinear curve fitting, one sample can “borrow information” from other genetically related samples during the estimation, preventing over-fitting to individual growth patterns resulting in a more robust model.

If we limit the expression of growth curve to specific functions, a one-step approach of the genomic prediction can be easily implemented to time-series phenotype data. It is known as the random regression models [16–18], in which functions that are linear combinations of features (e.g. Legendre polynomial) are used to express growth curve. The coefficients of the linear combinations are the parameters expressing representing growth curves, and are the response variables of the genomic prediction. This model structure enables the parameters to be estimated in one-step approach as one form of the mixed models [19]. Due to the easiness in implementation, the random regression model has been widely used for both animals [19] and plants [20, 21]. However, while the random regression models are good at prediction of time-series phenotype data, it is not suitable for interpretation of the growth characteristics using estimated parameters. To clarify the underlying genetic and environmental mechanisms determining growth curves, use of nonlinear functions in which roles of the parameters are well-known is more appropriate. Specifically, growth trajectories such as plant height often follow predictable patterns, unlike the fluctuating changes observed in animal body weight, where trajectory rules may not be as apparent. By utilizing well-known nonlinear functions, researchers can explore the genetic and physiological implications of trajectory shape parameters [6], moving beyond mere description of trajectory shape.

While this study focused on growth curves represented by nonlinear equations, crop growth in response to the environment has been described by a physiological model known as a crop growth model (CGM) [22–28]. In the context of the CGM, there has been some discussion about the advantages of one-step and two-step models [27, 28]. In the two-step model, CGM parameters are first estimated for each variety, and then models for specific genetic analysis, such as QTL analysis, genome-wide association study and genomic prediction, are applied to the estimated parameters. In the one-step model, the genetic effects on the variation of CGM parameters is modeled from the onset and CGM parameters for all varieties are estimated simultaneously. In the one-step model, Bayesian modeling is used due to its complexity, and the CGM parameters are estimated using computational algorithms such as Markov chain Monte Carlo (MCMC) [25], approximate Bayesian computation (ABC) [22, 24], and variational Bayes (VB) [23, 27]. Although our study focused on nonlinear growth curve models, genetic modeling and estimation algorithms for parameters have been studied in CGM as well.

Quantitative trait locus (QTL) analysis models have been developed using a one-step approach in which parameters of the nonlinear functions) are represented by a linear combination of locus effects [29, 30]. However, this model assumes that the effects of a single QTL are estimated and tested individually as fixed effects, and it cannot be used to make genomic predictions for traits that are controlled by multiple medium or small gene effects. In genomic selection, the effects of genome-wide markers are generally modeled as random effects [14, 31, 32]. Attempting to model the variation in the parameters of a nonlinear function with both random and fixed effects requires a framework that integrates mixed effects into a nonlinear function, that is, a nonlinear mixed-effects model (NLMEM). A few examples of NLMEMs have been applied to the genomic prediction of growth patterns [6, 33, 34], but they have not yet been fully explored.

NLMEMs are statistical models typically used to describe dynamic phenomena from repeated measurements of several individuals [35, 36]. They allow the simultaneous analysis of the data of several subjects in a single model by describing the variability of the phenomenon of interest in the studied population and the specificity of each individual at the same time. During many applications, such as in pharmacokinetics or in the study of growth curves, these models are generally utilized as models with random effects on the parameters. Statistical inferences in these models provide access to multiple pieces of information. Estimation of the mean values of the parameters and their variances allowed us to describe the phenomenon of global interest in the population. Some estimates of the individual parameters could also be obtained from these models, making prediction at the individual level as well as the comparison of the individual responses, for example, to select plants with high genetic abilities, possible. The advantage of NLMEMs is that all information can be inferred from the same statistical model. Studies on plant growth and genetic variation among cultivars are generally based on measuring the same phenotypic traits multiple times over time in plants of different cultivars. Therefore, it was appropriate to use mixed-effects models for this purpose. Although this approach was proposed several years ago in the field of animal genetics [12, 37], it is not widely used. One of the main objectives of this study was to illustrate the relevance of NLMEM in plant breeding when available data were repeated from several plants of different varieties.

In this study, NLMEM was applied to remotely sensed data of soybean growth during cultivation trials under different irrigation conditions. In the experiments used for the analysis, the whole-genome sequences of all the varieties were decoded, allowing their genetic relationships to be incorporated into the model. The influence of genetic relationships on the irrigation effects was also modeled. Specific tools are required to estimate and predict parameters from the NLMEM. Here, we used a framework that employs an algorithm derived from the Expectation-Maximization (EM) algorithm [38], that is, the stochastic Approximation EM (SAEM) algorithm [39], which is widely used in the NLMEM community and is very efficient in terms of computation time compared with those of standard MCMC methods. The effectiveness of NLMEM was examined by applying it to simulated data generated for a cultivation trial and observed data.

## 2 Materials and methods

### 2.1 Data description

A field trial was conducted from 2017 to 2022 in an experimental field with sandy soil at the Arid Land Research Center, Tottori University (35 °32 ^1^ N latitude, 134 °12^1^ E longitude, 14 m. above sea level). Soybean accessions registered in the National Agriculture and Food Research Organization Genebank (https://www.gene.affrc.go.jp/databases-core_collections_jw_en.php and https://www.gene.affrc.go.jp/databases-core_collections_wg_en.php) were used in this study. A total of 198 accessions consisted of 96 Japanese accessions, 96 world accessions from mini-core collections [40], and six additional accessions (GmWMC160, GmWMC157, GmWMC163, GmJMC002, GmJMC003, and GmJMC033). In 2017, 188 out of the 198 accessions were used for the analysis.

Whole-genome sequencing data of all 198 accessions [41] were used as previously described [13]. Biallelic SNPs with a minor allele frequency (MAF) ⩾ 0.025, missing rate < 0.05, and linkage disequilibrium < 0.95 were employed. Missing genotypes were imputed with Beagle 5.0 [42], using default parameter settings. The genome-wide SNP genotype data used in the analysis included 425,858 SNPs. Genotypes of individual alleles were scored as − 1 (homozygous for the reference allele), 1 (homozygous for the alternative allele), or 0 (heterozygous for the reference and alternative alleles).

Each plot contained four plants from the same variety. Sowing was performed at the beginning of July each year, followed by thinning after two weeks. Two watering treatment levels, control (C) and drought (D), were used to evaluate the genetic variations in the responses to water stress. Irrigation was applied daily for 5 h (7:009:00, 12:0014:00, and 16:0017:00), starting the day after the thinning in treatment C, while no water was applied in treatment D (see Figure 1). Plant growth was measured both manually and by remote sensing for some plants, which allowed the calibration of the UAV data. The R package missForest [43] was used to perform nonparametric missing value imputation using random forest. In the following section, we describe the analysis of UAV data collected in 2017. In 2017, there were up to 10 UAV measurements per plant (1-56 days after sowing). Weather data were used to convert the time units, expressed in terms of the number of days after sowing, into heat units. In summary, there are 752 plots of 188 genotypes, each with 425,858 markers, and phenotype data spanning 56 days. As measurements are aggregated by plot, leading to the monitoring of an average plant per plot, we will confuse the notions of plot and plant, and designate by plant the entity on which a single growth dynamic is observed.

**Fig 1.**
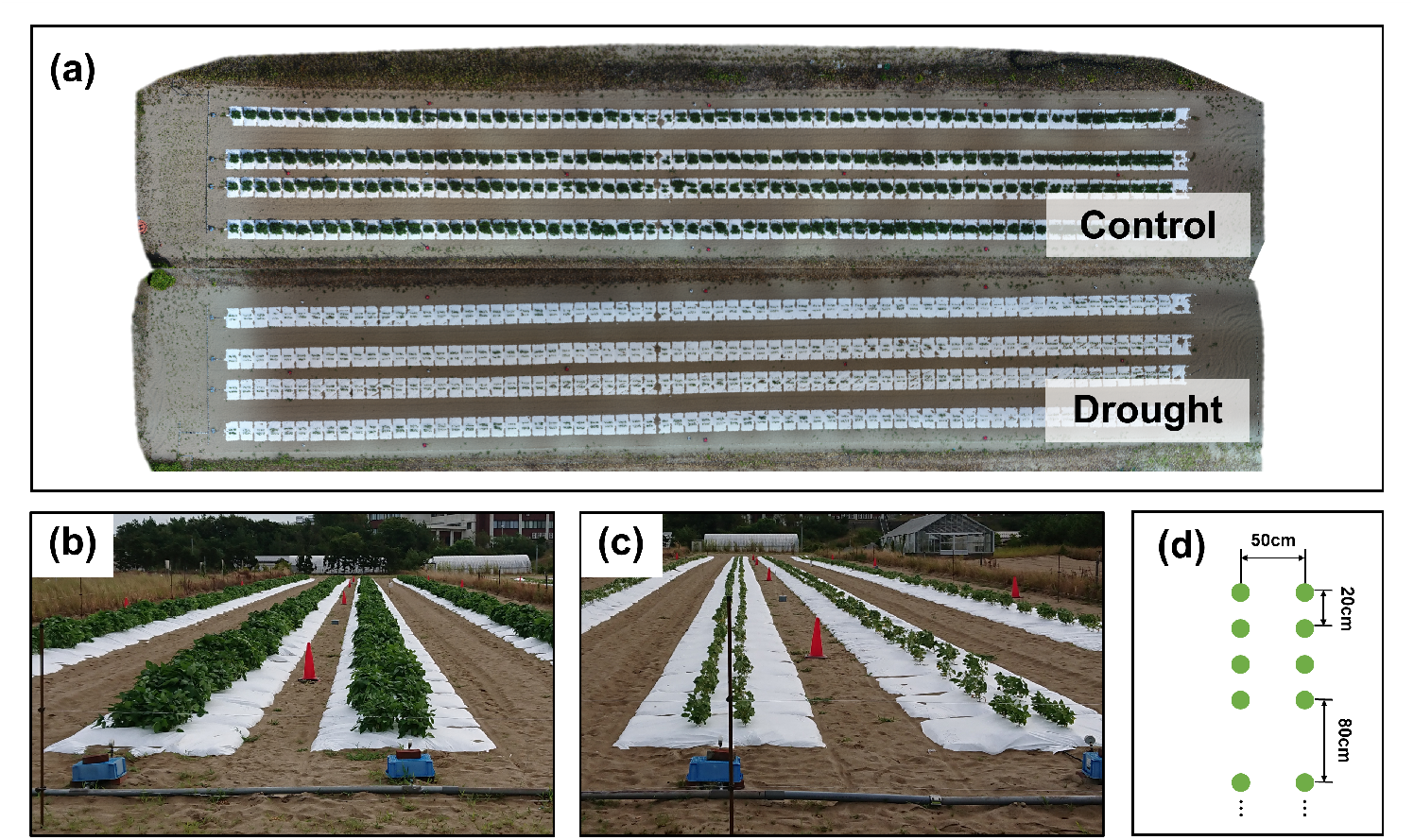
Illustration of the field design of the field experiment (Fig. 1 in [13]). The images are of the experiments in 2018, but their designs were the same. (a) An ortho-mosaic image of the field, obtained on August 25, 2018. The ortho-mosaic images were created for each treatment. (b,c) Ground level images of treatments C and D. (d) Planting pattern of plots made of 2 rows of 4 plants (green dots) and separated by 80 cm.

### 2.2 Modeling

We used NLMEMs with two levels of random effects to study genetic variability among soybean varieties. What sets this model apart from others in the literature is that it is here specified at the plant level rather than the variety level, thus making it possible to quantify the proportion of inter-plant variability due to covariates, the proportion of inter-plant variability due to genetics and the proportion of inter-plant variability unrelated to covariates and genetics. The general model is first described before presenting the model used in this study.

#### 2.2.1 Description of nonlinear mixed models with genetic effects

We assume *N*_*p*_ plants from *N*_*v*_ varieties *N*_*v*_ < *N*_*p*_ were observed. Let *n*_*i*_ be the number of observations for plant *I =* 1, …, *N*_*p*_, and *y*_*ij*_ ∈ **R** denote the *j*th observation of plant *i* obtained under the time condition *t*_*ij*_, *j* = 1, …, *n*_*i*_. The most generic way to formulate a NLMEM is as a hierarchical model, with each stratum describing a different level of variability in the data (see *e*.*g*., [35, 36]). As in [37], the following form is used:

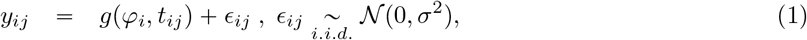

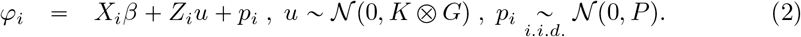

The first stratum, given by Equation (1), describes the data for each plant with the same statistical model but different parameters from one plant to another. *g*(*φ*_*i*_, *t*) is a known (nonlinear) function parameterized by *φ*_*i*_ ∈ **R**^*m*^, which describes the evolution of the growth process over time in an appropriate manner, with *m* the number of parameters that define each growth curve. Note that using the same *g* for all plants means that individual growths follow a common dynamic. Examples of *g*(*φ*_*i*_, *t*) can be found below. The second stratum, stated by Equation (2), aims at describing the differences between plants using covariates (*i*.*e*., explanatory measured variables characterizing the plants or their cultural conditions, arranged in *m*-by-*p* matrix *X*_*i*_ and *m*-by-*q*.*N*_*v*_ matrix *Z*_*i*_ with *q* the number of genetic effects per variety and *q*.*N*_*v*_ the product between *q* and *N*_*v*_, thus the total number of genetic values), genetic varietal effects 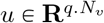 and plant-specific random effects *p*_*i*_ ∈ **R**^*m*^. More precisely, the term *X*_*i*_*β* describes the covariate effects common to all varieties using the fixed-effects vector *β* ∈ **R**^*p*^, whereas the term *Z*_*i*_*u* describes the covariate effects that differ from one variety to another using the genetic effect *u*. Matrices *X*_*i*_ and *Z*_*i*_ are the incidence matrices that relate plant *i* to the fixed effects and to the genetic values respectively.

Model (1)-(2) describes the inter-plant random variability of growth patterns *φ*_*i*_ by genetic variability with the *u* effects on the one hand and some variability that is not related to genetics with the *p*_*i*_ effects. The random varietal effect (*u*) and plant effect (*p*_*i*_) are assumed to be independent-centered Gaussian random variables whose respective variances are specified by means of the *q*-by-*q* covariance matrix *G*, the *N*_*v*_-by-*N*_*v*_ genetic relationship matrix *K* between the varieties, and the *m*-by-*m* covariance matrix *P* (see Equation (2)). Hence, the components in matrix *G* quantify the part of the variability between plants that is related to genetics, and the components in *P* quantify the variability between plants that is unrelated to covariates or genetics.

Note that this model is very flexible as it can be easily adapted according to the observed growth patterns simply by choosing an appropriate function *g* while still allowing an analysis of the genetic variability between varieties. It is important to note that the contribution of genetics to growth traits is contained in the genetic relationship matrix *K* and that the model does not include any effect of genetic markers in the fixed-effects part *X*_*i*_*β* as in [44].

To simplify the notation below, we denote by *θ* the set of unknown parameters in the model (1)-(2), composed of *β, P, G*, and *σ*^2^.

#### 2.2.2 Examples

We now use the same notations as above to introduce model examples that were used in the simulation study and the analysis of soybean experimental data.

##### The logistic growth curve

is a commonly used growth curve where function *g* is given by:

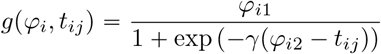

We considered a logistic growth curve with a constant and known scale parameter (*γ*), random asymptotic value (*φ*_*i*1_), and random time of half-maximum height (*φ*_*i*2_), hence the number of parameters that define each growth curve is *m* = 2 here. We considered no covariate effects or simple genetic effects on these parameters and thus defined *φ*_*i*_ = (*φ*_*i*1_, *φ*_*i*2_)^⊤^ as

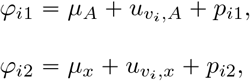

where *v*_*i*_ ∈ {1, …, *N*_*v*_} is the variety of plant *i*, vector *u*_*v*_ = (*u*_*v,A*_, *u*_*v,x*_)^⊤^ is the vector of genetic effects for variety *v* and 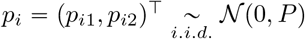. The vector *u* was obtained by the concatenation of the *N*_*v*_ genetic effect vectors *u*_*v*_’s such that *u* ∼ 𝒩 (0, *K* ⨂ *G*) with *G* the genetic covariance matrix and *K* the genetic relationship matrix between the *N*_*v*_ observed varieties. To connect with the general notations introduced in equations (1) and (2), the fixed effects vector in this model was *β* = (*μ*_*A*_, *μ*_*x*_)*T* and the covariate matrices are respectively 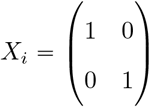 and the *N*_*v*_-block matrix 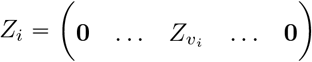 with 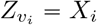. Matrices *P* and *G* were both 2 by 2 matrices.

##### The asymptotic regression curve

is another commonly used growth curve. Function *g* was defined as follows:

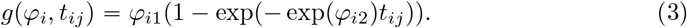

Asymptotic regression curves are defined with *m =* 2 parameters each. During the analysis of the soybean growth data presented in Section 3.2, we considered the cross-effect between genetics and irrigation conditions on both parameters *φ*_*i*1_ and *φ*_*i*2_ by defining *φ*_*i*_ *=* (*φ*_*i*1_, *φ*_*i*2_)^⊤^ as

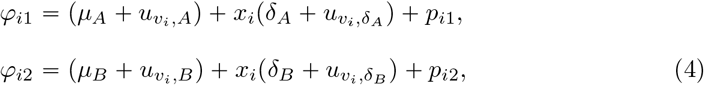

where *x*_*i*_ denotes the water condition plant *i* is subject to (*x*_*i*_ *=* 1 normal, *x*_*i*_ *=* 0 dry), *v*_*i*_ denotes the variety of plant *i*, and 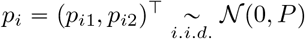. The vector of genetic effects for variety *v* ∈ {1, …, *N*_*v*_} was composed of four components: 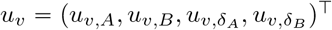. The vector *u* obtained by the concatenation of the *u*_*v*_’s is such that *u* ∼ 𝒩 (0, *K* ⨂ *G*) with *G* the genetic covariance matrix and *K* the genetic relationship matrix between the *N*_*v*_ observed varieties. Here, the fixed effects vector was *β =* (*μ*_*A*_, *μ*_*B*_, *δ*_*A*_, *δ*_*B*_)^*T*^ and the covariate matrices were 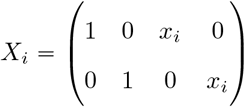 and the *N*_*v*_-block matrix 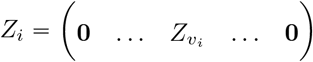 with 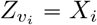. *P* and *G* were respectively a 2 by 2 and a 4 by 4 covariance matrices.

Remarks:

- Note that in both examples, we see from the equations defining *φ*_*i*1_ and *φ*_*i*2_ that the incidence matrix that relates plant *i* to the genetic values of the variety *v*_*i*_ it belongs to is the same as the incidence matrix that relates plant *i* to the fixed-effects. That is why *Z*_*vi*_ *= X*_*i*_ here, but this is not a constraint of our methodology, and we can easily consider models in which *Z*_*vi*_ and *X*_*i*_ are different.
- Note that there is no formal definition of heritability in these models. However, when a single genetic effect influences each growth trait, the *P* and *G* matrices have the same dimensions, and we can propose to evaluate the heritability of these traits using ratios calculated from their diagonal terms. In the example based on the logistic growth curve, this would give:

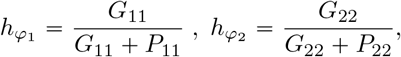

where *G*_*kk*_ and *P*_*kk*_ are the *k*-th diagonal terms of matrices *G* and *P* respectively, and 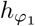 and 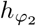 denote the heritabilities of growth traits related to growth parameters *φ*_*i*1_ and *φ*_*i*2_ respectively.

### 2.3 Algorithms

We intended to process the growth data by performing both a maximum likelihood estimation of the parameters and a maximum a posteriori estimation of the predicted genetic effects by using a unique statistical model for both tasks.

Model (1)-(2) is a nonlinear model with the random genetic and plant-specific effects *u* and *p*_*i*_ as latent variables. Because of these two features, neither the likelihood of observations nor the posterior density of genetic effects has an explicit form. The inferences in the model (1)-(2) fall within the scope of EM-like algorithms. Here, owing to the model nonlinearity, we propose to tackle both parameter estimation and genetic effects prediction using the SAEM algorithm [39] instead of the EM algorithm.

#### 2.3.1 General overview of the SAEM algorithm

The EM algorithm [38] is an iterative algorithm in which the estimate of an unknown parameter is updated by maximizing the expectation of the log-likelihood of the complete data (*i*.*e*., observed and latent data) conditional on the observations. Considering a general latent variable model with *y* and *z* as the observed and the unobserved variables respectively and unknown parameter *θ*, iteration [*k*] of the EM algorithm requires the computation of **E**[log *p*(*y, z*; *θ*)|*y, θ*^[*k*−1]^] with *θ*^[*k* − 1]^ the estimation of *θ* obtained at the previous iteration [*k* − 1]. When **E**[log *p*(*y, z*; *θ*)|*y, θ*^[*k*−1]^] is not available in a closed form, the SAEM algorithm replaces its exact calculation with a stochastic approximation as follows:

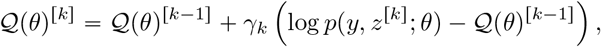

where *z*^[*k*]^ is a random draw from the conditional distribution *p*(*z*|*y, θ*^[*k*−1]^], either directly or using Monte Carlo methods, if the exact simulation under *p*(*z*|*y, θ*^[*k*−1]^] is unfeasible [45]. (*γ*_*k*_)_*k*>0_ is a sequence of positive decreasing step sizes, such that *γ*_1_ = 1, *∑γ*_*k*_ *=* ∞ and 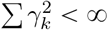. Hence, the calculation of the complete log-likelihood and simulation of the latent data are two central points in the implementation of the algorithm.

#### 2.3.2 Parameter estimation

Within the model (1)-(2), the observations are the available height measures from the *N*_*p*_ plants, 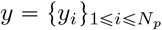 where 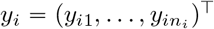 is the vector of observations for plant *i*, and the unknown parameter *θ* is composed of the fixed-effects *β*, the residual variance *σ*^2^ and the covariance matrices *G* and *P*. As suggested in [37], a possible and convenient choice for the missing data is *z =* (*φ, u*), where 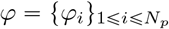 is the set of plant growth parameters and *u* is the vector of genetic effects. Therefore, the likelihood of the complete data can be decomposed into:

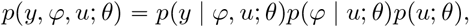

where each element of the right-hand term represents an easily computable Gaussian density. An exact simulation under *p*(*φ, u*|*y*; *θ*) is not feasible, but the Metropolis within Gibbs [46] is easy to implement by exploiting the hierarchical dependency between *y, φ*, and *u* and using that the exact simulation under *p*(*u*|*φ*; *θ*) is feasible. Iteration [*k*] of the algorithm can thus be implemented as follows:

- *Simulation step:*
  – Draw *φ*^[*k*]^ under 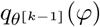 a transition kernel of a Monte Carlo Markov Chain algorithm targeting *p*(*φ*|*u*^[*k*−1]^, *y, θ*^[*k*−1]^) as a stationary distribution
  – Draw *u*^*[k]*^ under *p*(*u*|*φ*^*[k]*^, *θ*^[*k*−1]^)
- *Stochastic Approximation step:* compute

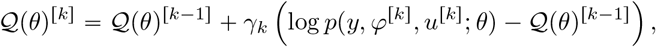

where *γ*_1_ = 1, ∑ *γ*_*k*_ *=* ∞ and 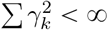.
- *Maximization step:* compute

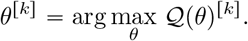

The succession of these three steps is repeated until the algorithm convergence. Implementation details and justifications are provided in Appendix S1 (see Algorithm 2).

#### 2.3.3 Prediction of the varietal genetic effects

The varietal genetic effects can be predicted by computing

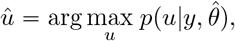

where 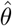 denotes the estimated parameter value obtained at the convergence of the algorithm described in the previous section. The conditional density *p*(*u*|*y, θ*) has no explicit form in model (1)-(2), but as *p*(*u*|*y, θ*) = ∫*p*(*u*|*φ*; *θ*)*p*(*φ*|*y, θ*)*dφ*, it is possible to use the SAEM algorithm again by considering *φ* as the latent variable and *u* as the element to be estimated. Iteration [*k*] of the obtained algorithm is decomposed as follows:

- *Simulation step:* draw *φ*^*[k]*^ under 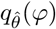 a transition kernel of a Monte Carlo Markov Chain algorithm targeting 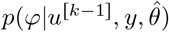 as a stationary distribution
- *Stochastic Approximation step:* compute

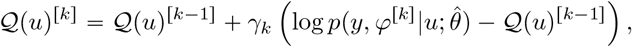

where *γ*_1_ = 1, ∑ *γ*_*k*_ *=* ∞ and 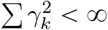.
- *Maximization step:* compute

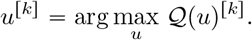

Implementation details are provided in the Appendix (see Algorithm 2).

### 2.4 Numerical study design

Our study is in two parts. The first focuses on the quality of the model’s parameter estimates, which is crucial for accurately understanding and characterizing general plant behavior and the heritability of growth patterns. The second looks at predictive performance, exploring different scenarios.

#### 2.4.1 Parameter estimation

We used the two-parameter logistic growth model introduced in section 2.2.2 with the following parameter values: *μ*_*A*_ *=* 50, *μ*_*B*_ *=* 13, 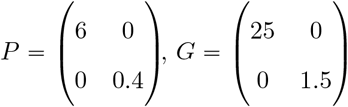, and *σ*^2^ = 4. Growth data are simulated on time interval r1, 45s according to different scenarios to illustrate the performance of our approach based on nonlinear mixed-effects models and the SAEM algorithm in terms of parameter estimation, and to make recommendations for data collection.

1. *Scenario 1* is used to study the effect of the number of observed varieties and of the number of plants per variety on the performances of our methodology. The total number of plants is *N*_*p*_ *=* 2000, and the number of observed varieties is either *N*_*v*_ *=* 50 or *N*_*v*_ *=* 100. We consider a high frequency of data collection over the curves by simulating one data point at each unit time step leading to 45 observations per plant.
2. *Scenario 2* is used to study the effect of sampling frequency along the curves. We use for *N*_*v*_ and *N*_*p*_ a similar value as for the real data, *N*_*p*_ *=* 800 and *N*_*v*_ *=* 200. We simulate regularly spaced observations along the plant growth curves, with a total of either 45 (rich design) or 10 (sparse design) observations per curve.

50 datasets are simulated for each scenario and for each combination of *N*_*p*_, *N*_*v*_, and sampling frequency along the curves. The SAEM algorithm is applied to each of them to estimate the *θ* parameters (*i*.*e. β, σ*^2^, *G* and *P*). The algorithm is run for 1500 iterations in total, with step sizes for the stochastic approximation defined as *γ*_*k*_ *=* 1 if *k* < 1300 and *γ*_*k*_ *=* (*k* − 1300)^−2/3^ otherwise. This setting corresponds to defining a burn-in phase of 1300 iterations aimed at converging quickly to the neighborhood of the maximum likelihood estimator, followed by a refining phase in which the estimation is smoothed by decreasing the step sizes. Note that we use here a large number of iterations to guarantee good convergence of the algorithms on all simulated datasets; however, in practice, we observe that the convergence is reached far before the end of the 1500 iterations.

#### 2.4.2 Prediction

Once the parameters have been estimated, it is possible to estimate genetic effects *û* and derive predictions of growth traits and observations at any time point *t* for a given plant *i* using the following formulas :

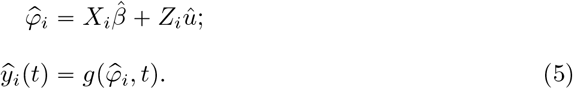

Depending on the objectives of the study, both types of prediction, 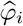 and 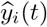, may be of interest. The logistic growth model introduced in section 2.2.2 is also used to evaluate the prediction accuracy of our methodology, but this time including a covariate *x*_*i*_ in the definition of the growth parameters acting as a proxy of the environment:

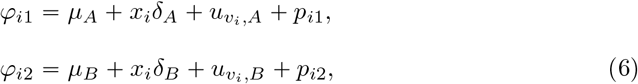

where *v*_*i*_ denotes the variety of plant 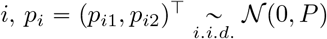 and *u* is obtained by the concatenation of the *u*_*v*_’s is such that *u* ∼ 𝒩 (0, *K* ⨂ *G*). The following parameter values are used: *μ*_*A*_ *=* 50, *μ*_*B*_ *=* 25, *δ*_*B*_ *=* 25, *σ*^2^ = 8, *P =* diag(3, 0.1) and *G* is defined as *G = kP* with *k* ∈ {1, 2} in order to explore different heritability levels (50% and 67% respectively). Predictive performances of our methodology are evaluated under different scenarios suggested by the literature which we name cv0, cv1, and cv2 as in other articles (see *e*.*g*. [47]). In each scenario, and for each level of heritability, we simulate 50 datasets according to model (6) each divided into a training set and a validation set. The original datasets contain *N*_*v*_ *=* 225 varieties each, six plants per variety are simulated, two in each environment, leading to a total number of plants *N =* 1350. Parameter and genetic effects are estimated from the training set using the SAEM algorithms, and the values of 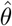 and *û* are used in formulas (5) to compute growth parameters and observations predictions in the validation set. Scenario cv0 is designed to evaluate predictions on new environmental conditions. Training and validation samples are made up by leaving one environment out and predictions are derived for each variety in the left-out environment. In this scenario, every variety is observed in the training set but in environments different from the one to be predicted. Scenario cv1 is designed to evaluate predictions on new varieties in any environment. Varieties present in the validation set are not observed in the training set. The kinship matrix is used to deduce the genetic effects of these new varieties from those estimated for the varieties in the training set, and then formula (5) is applied. Training and validation sample sizes are defined as 90% and 10% of the initial data set, respectively. Scenario cv2 is designed to evaluate predictions on varieties not evaluated in one environment, but evaluated across other different environments. Training and validation samples are made up by leaving some combinations between varieties and environments out. In this scenario, all varieties and environments are in the training set so that the estimated genetic effects can directly be used in formula (5) to compute predictions. As above, training and validation sample sizes are defined as 90% and 10% of the initial data set, respectively. To evaluate the three scenarios and in particular scenario cv0 on the same simulated data sets, *x*_*i*_ is quantitative and observations under values 1, 2, 3 for *x*_*i*_ are simulated.

## 3 Results

### 3.1 Simulations

#### 3.1.1 Parameter estimation

The simulation results related to parameter estimation are depicted in Fig 2, 3, 4 for Scenario 1, and on Fig 5, 6, 7 for Scenario 2. We see that all the parameters are correctly estimated, *i*.*e*., with very limited bias, regardless of the number of varieties, plants per variety, and time points per curve used for the data simulation.

**Fig 2.**
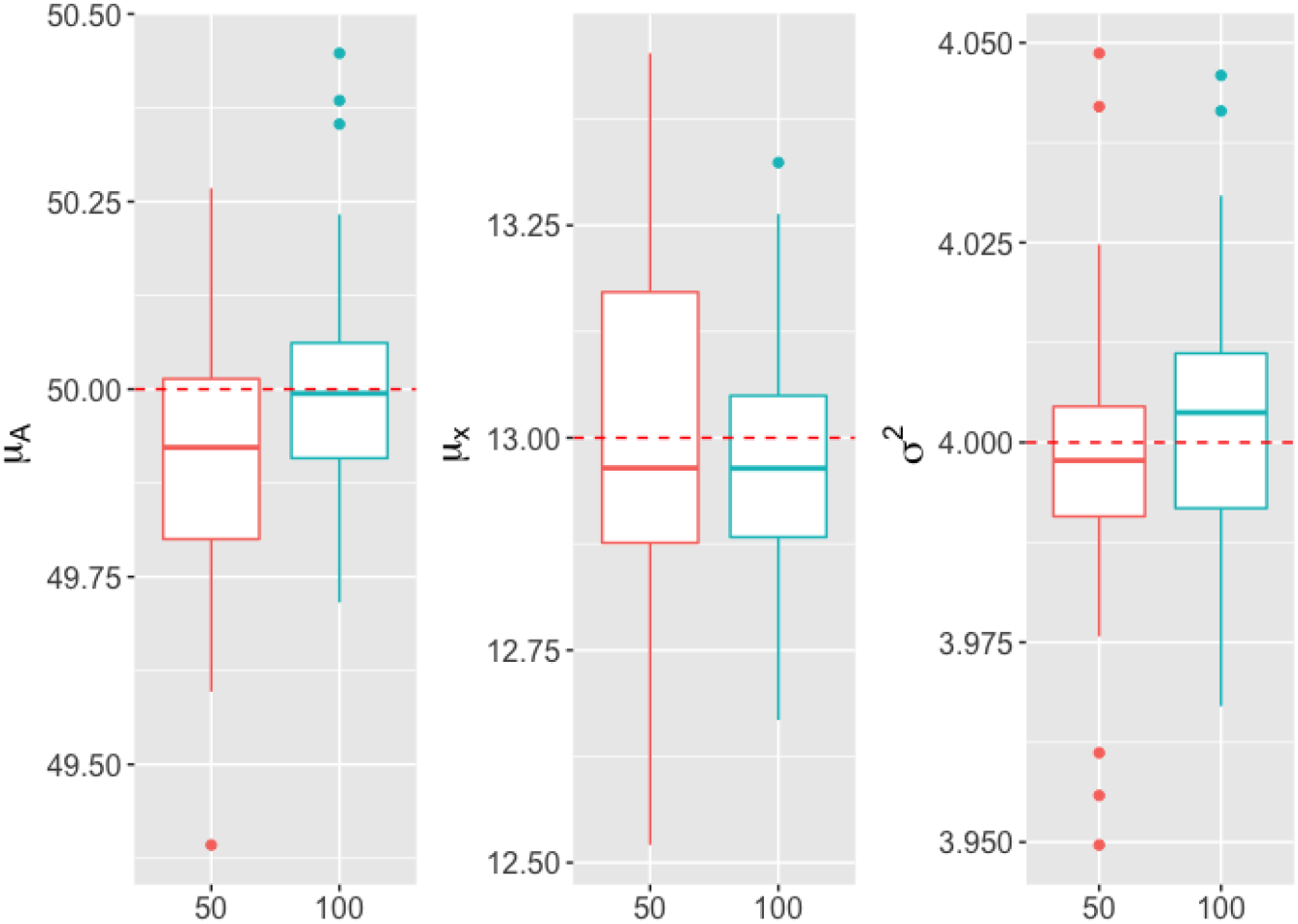
Boxplots of parameters *μ*_*A*_, *μ*_*x*_ and *σ*^2^’s estimations (y-axis) computed from 50 simulated datasets including different numbers of varieties (x-axis). Red-colored boxplots correspond to *N*_*v*_ = 50 varieties, blue-colored boxplots correspond to *N*_*v*_ = 100 varieties, and the red dotted line indicates the true parameter value.

**Fig 3.**
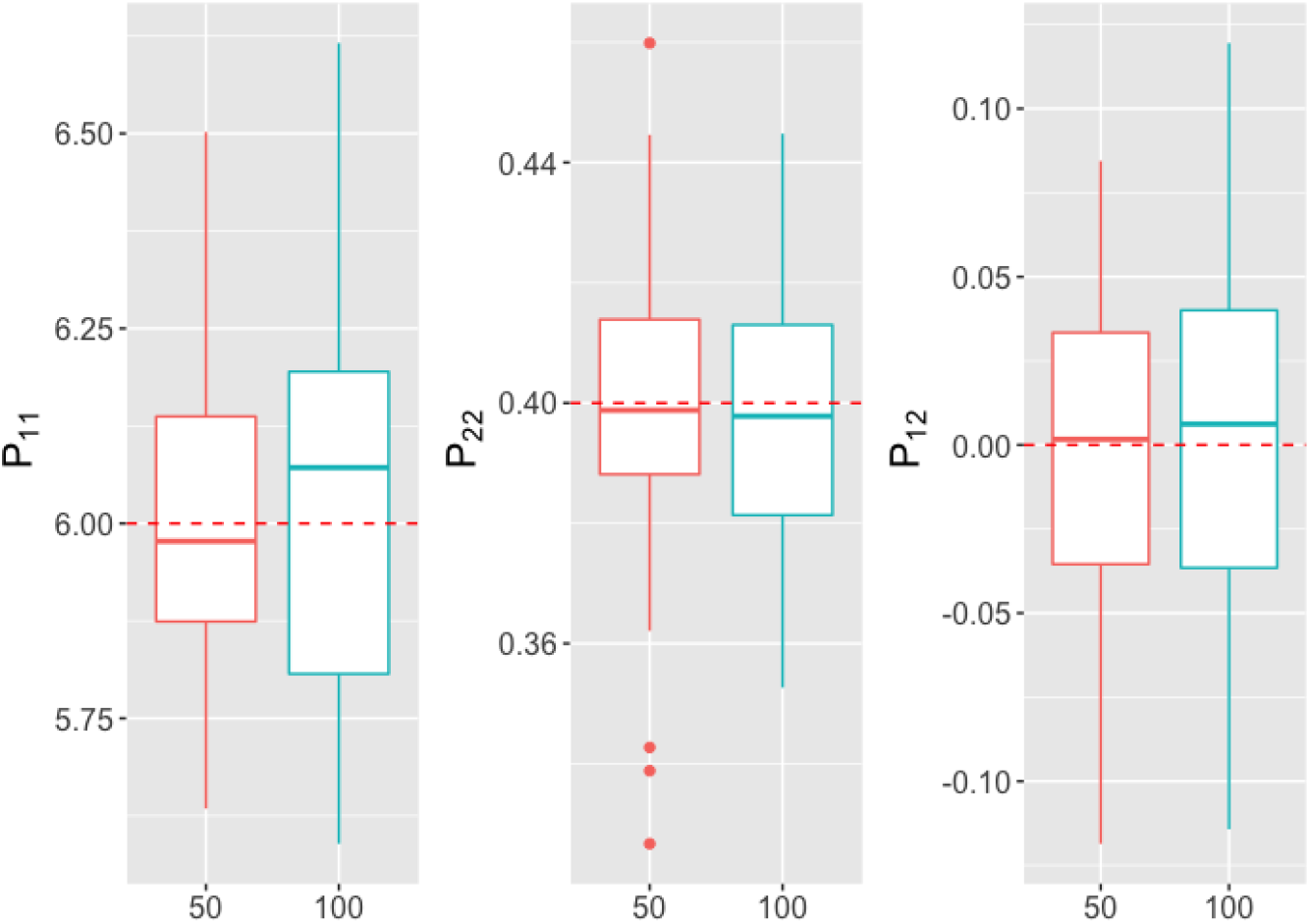
Boxplots of *P* variance and covariance parameters’ estimations (y-axis) computed from 50 simulated datasets including different numbers of varieties (x-axis). Red-colored boxplots correspond to *N*_*v*_ *=* 50 varieties, blue-colored boxplots correspond to *N*_*v*_ *=* 100 varieties, and the red dotted line indicates the true parameter value.

**Fig 4.**
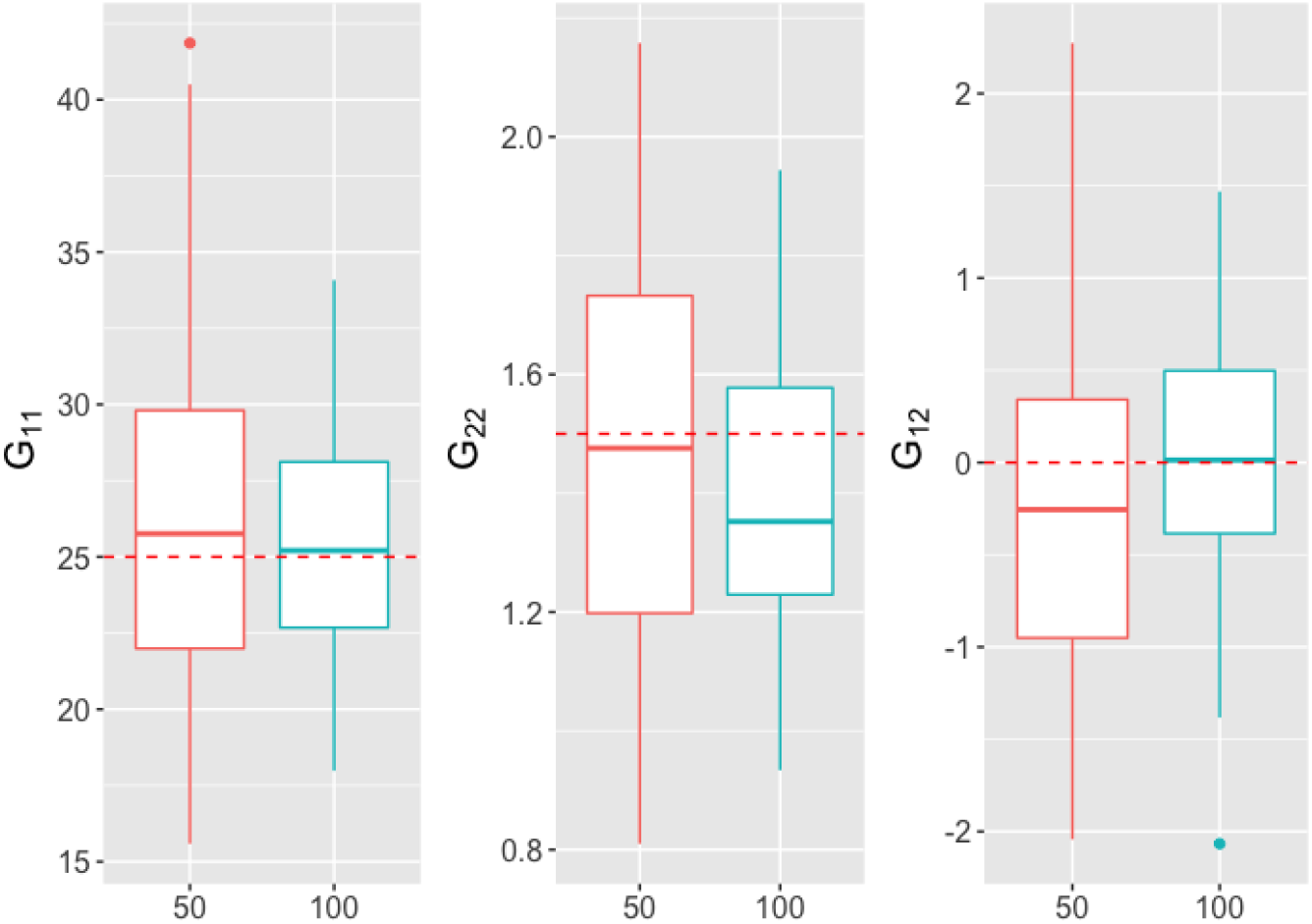
Boxplots of *G* variance and covariance parameters’ estimations (y-axis) computed from 50 simulated datasets including different numbers of varieties (x-axis). Red-colored boxplots correspond to *N*_*v*_ *=* 50 varieties, blue-colored boxplots correspond to *N*_*v*_ *=* 100 varieties, and the red dotted line indicates the true parameter value.

**Fig 5.**
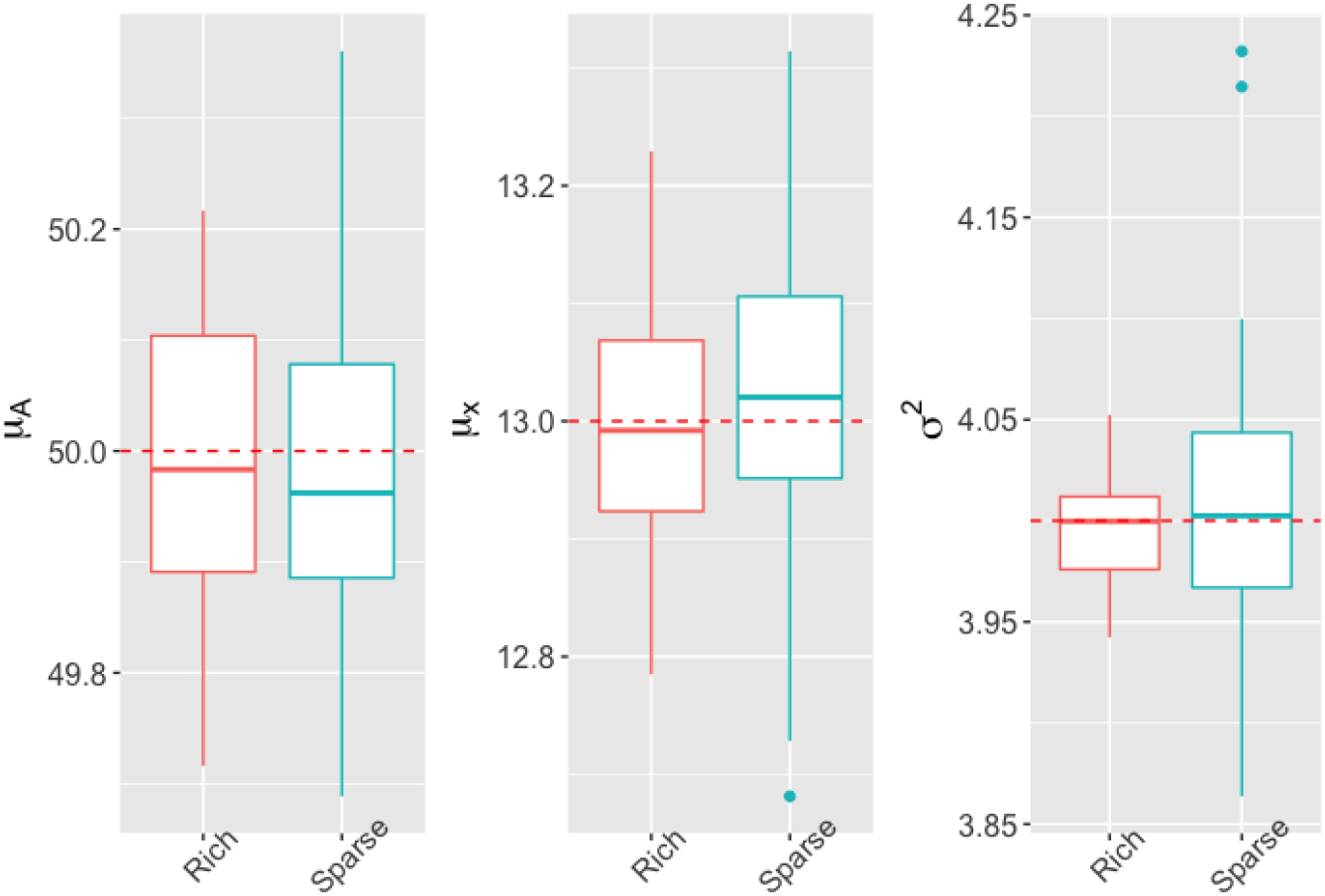
Boxplots of parameters *μ*_*A*_, *μ*_*x*_ and *σ*^2^’s estimations (y-axis) computed from 50 simulated datasets including different numbers of observations per plant (x-axis). Red-colored boxplots correspond to a rich design with 45 observations per plant, blue-colored boxplots correspond to sparse design with 10 observations per plant, and the red dotted line indicates the true parameter value.

**Fig 6.**
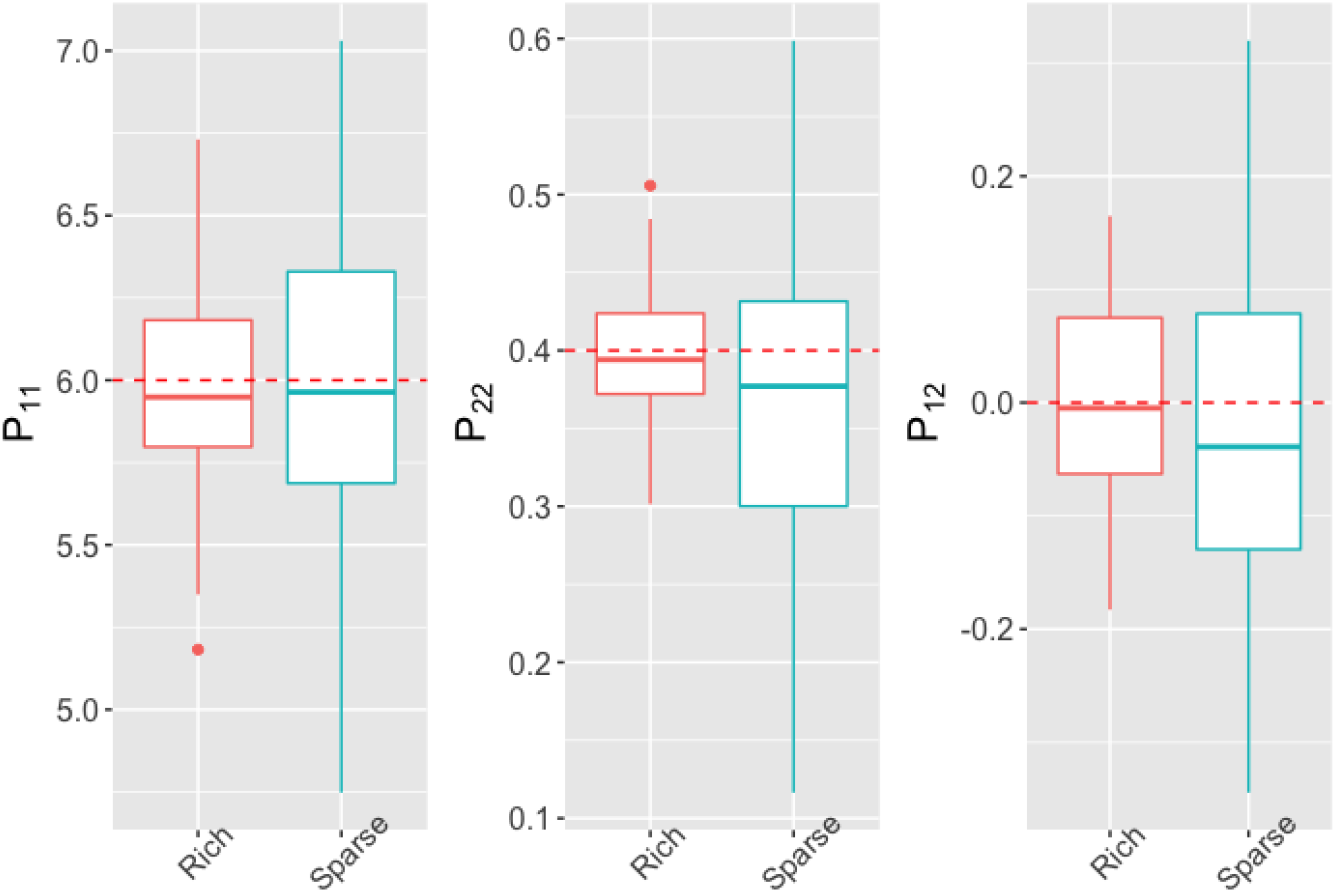
Boxplots of *P* variance and covariance parameters’ estimations (y-axis) computed from 50 simulated datasets including different numbers of observations per plant (x-axis). Red-colored boxplots correspond to a rich design with 45 observations per plant, blue-colored boxplots correspond to sparse design with 10 observations per plant, and the red dotted line indicates the true parameter value.

**Fig 7.**
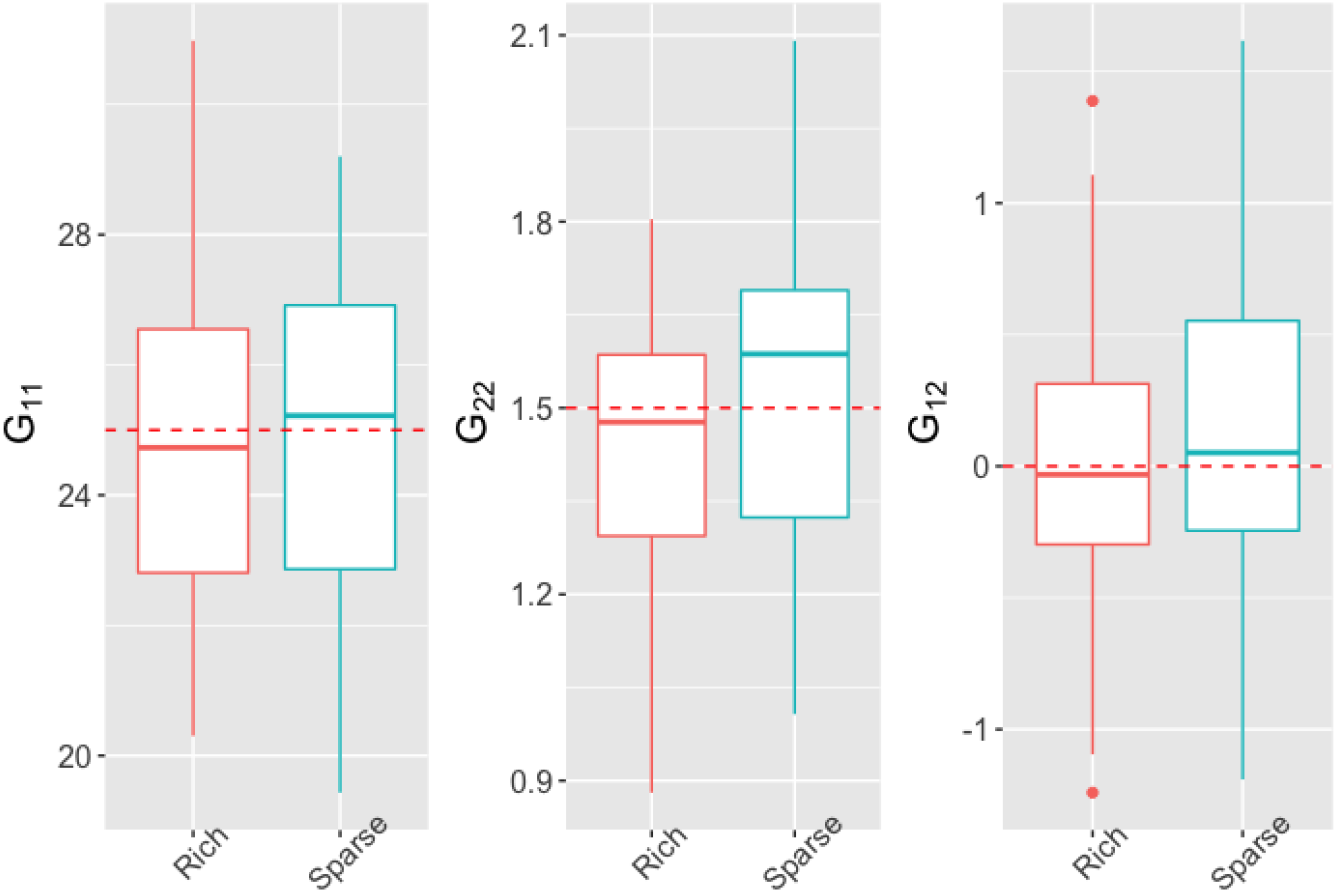
Boxplots of *P* variance and covariance parameters’ estimations (y-axis) computed from 50 simulated datasets including different numbers of observations per plant (x-axis). Red-colored boxplots correspond to a rich design with 45 observations per plant, blue-colored boxplots correspond to sparse design with 10 observations per plant, and the red dotted line indicates the true parameter value.

From Fig 2, 3, and 4, it appears that for the values considered for the number of varieties considered in Scenario 1, changing *N*_*v*_ primarily affects the variance of the estimates. Specifically, increasing *N*_*v*_ increases the precision of the intercept and genetic variance estimates. However, increasing *N*_*v*_ reduces the precision of the estimate of the covariance matrix *P*. The rationale for this behavior could be that, because this experiment is conducted on a fixed total number of simulated plants (*N*_*p*_), increasing *N*_*v*_ resulted in a decrease in the number of observed plants per variety. Increasing the number of observed plants per variety improved the quality of the estimations of *P*.

Fig 5, 6, and 7 show that for simulated data with similar values of *N*_*v*_ and *N*_*p*_ to the real data analyzed in the next section, the model parameters are also estimated with very limited bias. For these values of *N*_*v*_ and *N*_*p*_, decreasing the number of observations per curve does not bias the estimates, but increases their variance. Thus, the experiments conducted under the second simulation scenario show that it is possible to estimate satisfactorily the parameters of an NLMEM of the form (1)-(2) under sampling conditions similar to those of the real data collected in Tottori in 2017.

#### 3.1.2 Prediction

The prediction accuracies of both varietal growth parameters and height measures are examined by using relative differences between predicted and simulated values 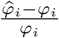 and 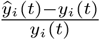. The results are depicted in Figures 8 and 9. We note from Figure 9 lower-quality predictions for the first measurement times, due to a lower signal-to-noise ratio than for other measurement times. Overall, the predictions of the parameters and observations at each time are slightly worse in scenario cv1 than in scenarios cv0 and cv2, which is an expected result, but satisfactory overall. Despite this, predictive performance is very similar between scenarios cv0, cv1 and cv2, and appears to be little influenced by heritability in the value ranges explored.

**Fig 8.**
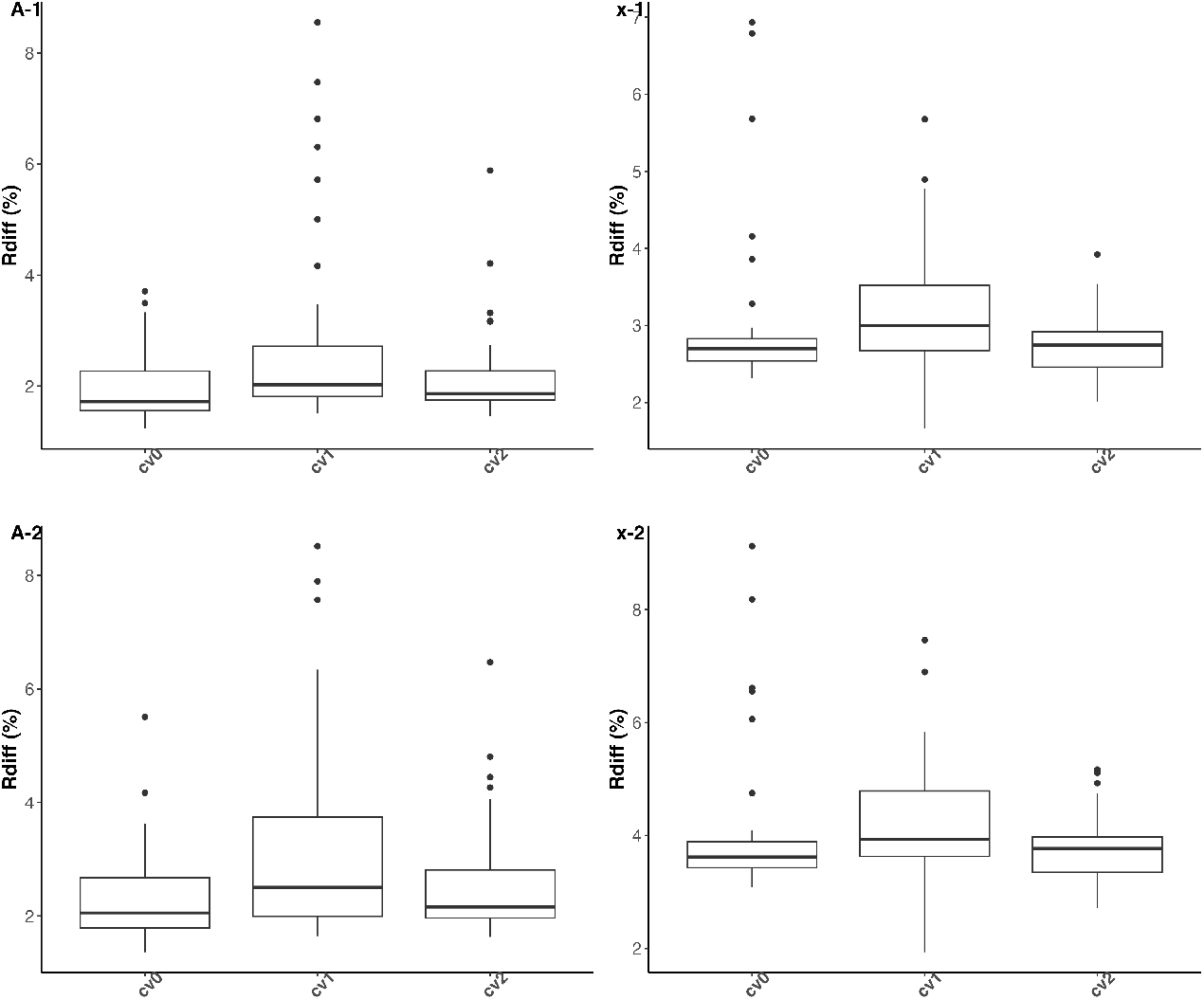
Boxplots of relative differences between predicted and simulated values of parameters A (left column) and x (right column) of the logistic growth curve under prediction scenarios cv0, cv1 and cv2 (x-axis). Panels (A-1) and (x-1) (resp. (A-2) and (x-2)) represent the results obtained with an heritability level of 50 % (resp. 67%) on each growth parameter.

**Fig 9.**
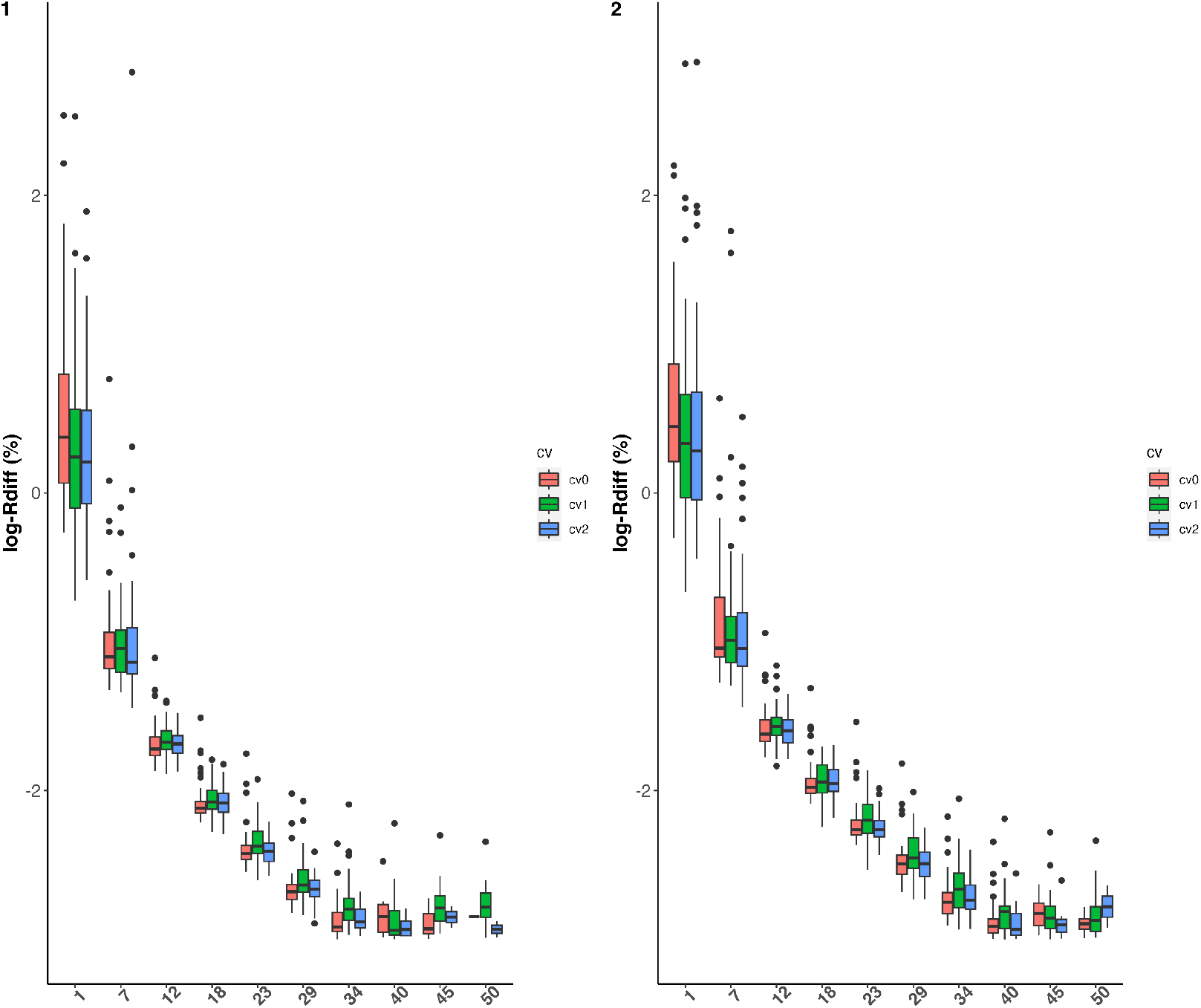
Boxplots of relative differences between predicted and simulated values of height measures (y-axis) per time point (x-axis) on a log-scale. Prediction scenarios cv0, cv1 and cv2 are represented by different colors. Left (resp. right) panel represents the results obtained with an heritability level of 50 % (resp. 67%) on each growth parameter.

### 3.2 Real data analysis

#### 3.2.1 Analysis based on the NLMEM approach

The dataset was composed of *N*_*p*_ *=* 752 plants of *N*_*v*_ *=* 188 varieties observed at 10 time-points. Some data transformation was required before applying the NLMEM approach to the 2017 Tottori UAV data. The growth dynamics represented in Fig 10 correspond to that of logistic growth, but the observations show strong heteroscedasticity over time. To avoid strong numerical instabilities, the logarithms of the measured heights rather than on the heights themselves were used. The log growth curves are represented in Fig 11. Their shape suggests the use of an asymptotic regression curve for their analysis, as defined in Equation (3).

**Fig 10.**
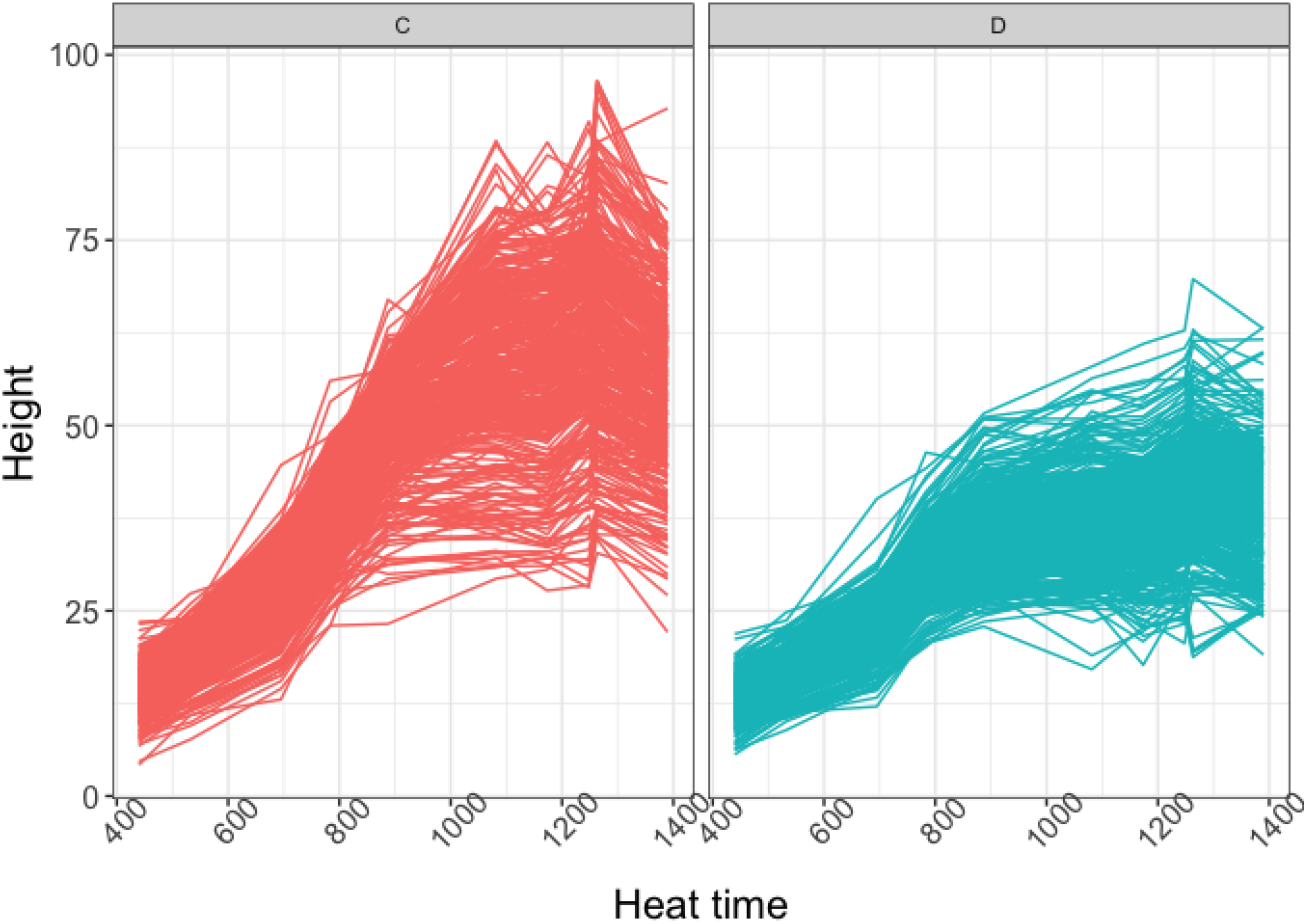
2017 Tottori UAV data by water condition. Y and x-axis respectively represent the height of the plants and heat time points. Panel C (left-red): normal water condition, Panel D (right-blue): dry condition.

**Fig 11.**
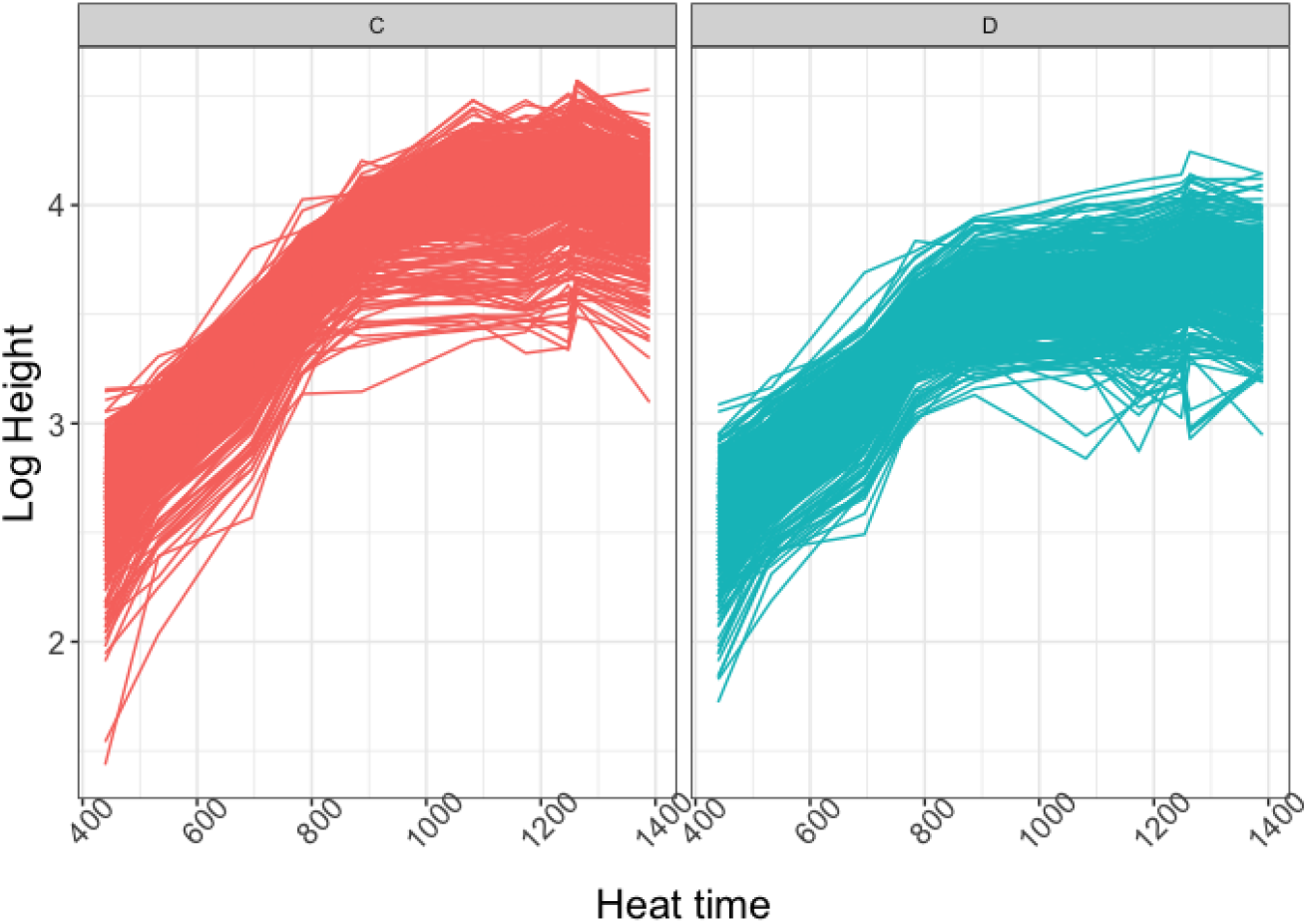
2017 Tottori UAV log-transformed data by water condition. Y and x-axis respectively represent the logarithm of the height of the plants and heat time points. Panel C (left-red): normal water condition, Panel D (right-blue): dry condition.

In the growth model, because it is applied to the logarithm of height measures, *φ*_*i*1_ represents the maximum expected log height of the plant, and exp(−*φ*_*i*2_) is the growth rate. The effects of irrigation conditions and genotypes (varietal differences) on the growth mechanisms were studied using model (4) on (*φ*_*i*1_, *φ*_*i*2_). In this model, *μ*_*A*_ and *μ*_*B*_ are the averages of *φ*_*i*1_ and *φ*_*i*2_, respectively, under drought conditions, and *δ*_*A*_ and *δ*_*B*_ are the average effects of irrigation on these two quantities, respectively. As mentioned in Section 2.2.2, the genetic effects of each variety in model (4) have four components: *u*_*v*_ *=* (*u*_*v,A*_, *u*_*v,B*_, *u*_*v,δ*_*A, u*_*v,δ*_*B*)^⊤^, which measures the specificity of each variety with respect to the logarithm of the maximum height and growth rate under drought conditions as well as the effects of irrigation. *G* is the covariance matrix of genetic effect *u*_*v*_.

The SAEM algorithm for parameter estimation was run for 400 iterations, of which 300 was run for burn-in with step sizes defined as *γ*_*k*_ *=* (*k* − 300)^−2/3^ in the refining phase. The algorithm for computing the genetic effects was run on 1000 iterations of which 800 for burn-in and step sizes defined as *γ*_*k*_ *=* (*k* − 800)^−2/3^ after burn-in. Sample convergence graphs of the SAEM algorithm for both parameter estimation and prediction are provided in Appendix S2. Both the algorithms converged satisfactorily.

We obtained the following parameter estimates:

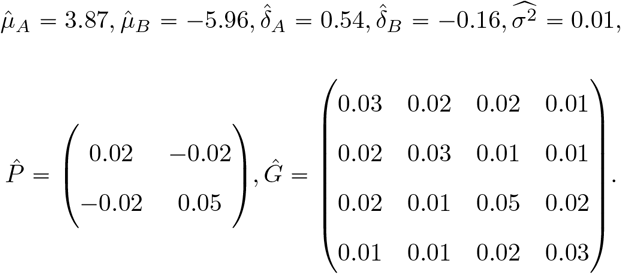

The estimates suggest that the soybean log-growth curves reached a plateau with a value of 3.87 on average under drought conditions and 4.41 under normal irrigation conditions. The growth rate parameter *φ*_*i*2_ had an average value of −5.96 among soybean varieties under drought conditions and −6.12 under normal irrigation conditions, corresponding to slower growth under drought conditions. These results are consistent with what is observed in Fig 11.

The estimations of matrices *P* and *G* provide a quantification of the different sources of variability in growth patterns and allow for the evaluation of genetic and marginal correlations between varietal effects or between growth parameters. The genetic correlation matrix, obtained by converting Ĝ into a correlation matrix, is estimated to

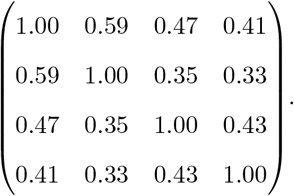

This allowed us to assess the correlations between genetic effects of any variety All genetic correlations in the above matrix are positive, implying that an increase in the genetic effect on the maximum height under drought conditions *φ*_*i*1_ would be associated with an increase in the genetic effect on *φ*_*i*2_, and thus on the growth rate, and a more pronounced genetic effect on the sensitivity to irrigation variations.

Correlations between the growth patterns *φ*_*i*1_ and *φ*_*i*2_, rather than the genetic effects, were also investigated. According to Equation (2) used in our modeling approach, the marginal covariance matrix of *φ*_*i*_ *=* (*φ*_*i*1_, *φ*_*i*2_)^⊤^ is

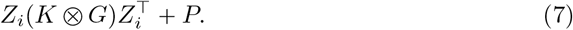

This equation includes the contribution of two parts: one that is exclusively due to genetics given by 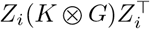, and the other *P* due to uncontrolled environmental conditions and random plant-specificity. Notably, because of the genetic relationship matrix *K* in the above formula, the covariance of *φ*_*i*_ changes slightly from one variety to the other, in addition to its variation between the irrigation conditions because of the changes in *Z*_*i*_ (*i*.*e*., drought and normal irrigation). Using formula (7) and the estimated values 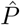 and *Ĝ*, by averaging over the *N*_*v*_ *=* 188 varieties and then converting them into correlation matrices, we found that the average genetic correlation between *φ*_*i*1_ and *φ*_*i*2_ is 0.002 under normal irrigation conditions and 0.04 under drought conditions, whereas the average marginal correlations are −0.18 and −0.25 under normal and drought conditions, respectively. This difference in sign between the genetic and marginal correlations suggests a stronger effect of the environment than that of genetics on the growth behavior of soybeans. The respective contributions of genetics and the environment to the variance of each parameter were also quantified using heritability values. The heritability of each of the growth curve parameters *i*.*e*., the maximum log-height (*φ*_*i*1_) and the growth rate (*φ*_*i*2_) under drought conditions, denoted by *h*_*A*_ and *h*_*B*_ respectively, and the heritability of the irrigation effect on each of these parameters, denoted by 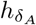 and 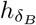, can be deduced by calculating the following ratios

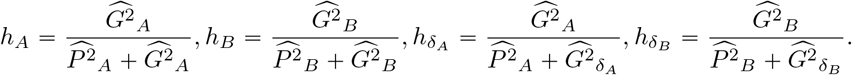

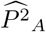 and 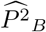 are the diagonal terms of 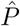, and 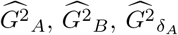, and 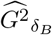 are diagonal terms of *Ĝ*. We obtained the following values for heritability 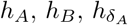, and 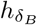 : 52%, 66%, 36%, and 41%, respectively. The maximum plant height and growth rate under the reference conditions showed more than half of the variability explained by genetics. This justifies considering the prediction of varietal effects on these parameters for potential varietal selection. In contrast, the contribution of genetics to the effects of irrigation on growth patterns,*i*.*e*., the contribution of genetic and irrigation interactions, appears to be relatively small; thus, genomic selection based on these effects would be less accurate for this soybean population. Broadly speaking, the predicted parameters 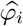 under the two conditions could be used in practice to select varieties that perform better (*e*.*g*., those with larger plant size or faster growth rate) under drought conditions or are less sensitive to climate change. The predicted varietal growth parameters are shown in Fig 12, by the water condition or by crossing the normal and drought conditions. The color of the dots represents the type of variety as defined in [41] (“Japan”, “World” and “Primitive”). Overall, the Japanese varieties had a lower growth rate than those of the “Word” and “Primitive” varieties under both conditions. The “Primitive” varieties grew faster in both conditions, attaining high maximum sizes among all the varieties. The “World” varieties had intermediate growth rates and reach the highest heights.

**Fig 12.**
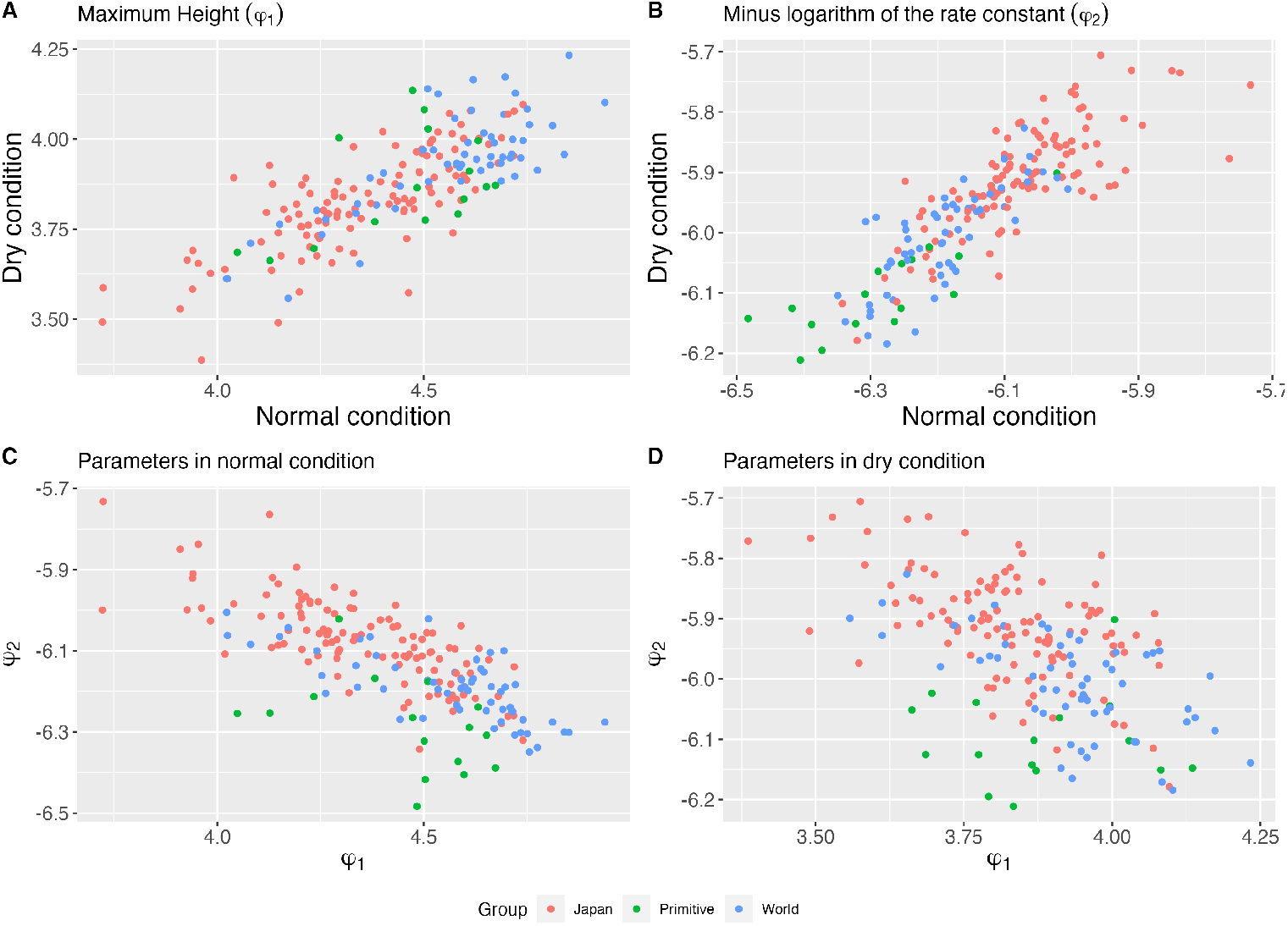
Varietal predicted parameters under both water conditions per groups of varieties. Panel A: Predicted values of growth parameter *φ*_1_ in drought condition (y-axis) as a function of predicted values of growth parameter *φ*_1_ in normal condition (x-axis); Panel B: Predicted values of growth parameter *φ*_2_ in drought condition (y-axis) as a function of predicted values of growth parameter *φ*_2_ in normal condition (x-axis); Panel C: Predicted values of growth parameter *φ*_2_ in normal condition (y-axis) as a function of predicted values of growth parameter *φ*_1_ in normal condition (x-axis); Panel D: Predicted values of growth parameter *φ*_2_ in drought condition (y-axis) as a function of predicted values of growth parameter *φ*_1_ in drought condition (x-axis). Colors correspond to different groups of varieties (red: Japan, green: Primitive, blue: World).

The robustness of predictions derived from the estimated model was assessed by running 5-fold cross-validation four times under scenarios cv1 and cv2 described in Section 2.4.2, allowing 20 repetitions in each scenario. Cross-validation under scenario cv0 was not possible here because the only covariate providing information on the environment is the water condition, which is qualitative. Training and validation samples under scenario cv1 (resp. cv2) are made up by randomly dividing varieties (resp. combinations between varieties and hydric condition) into 5 groups and then leaving all the observations of varieties (resp. combinations between varieties and hydric condition) in the same group, one group at a time. Parameters *θ* and genetic effects *u* are estimated by using the training sample, from which predictions of height measures are deducted for varieties or combinations between varieties and hydric conditions composing the validation sample according to the steps detailed in Section 2.4.2. Relative differences between predicted and observed values are computed and depicted in Figure 13. Smaller relative differences are a sign of better quality prediction. We see from Figure 13 that the predictive performance of the model is satisfactory overall, with relative differences between observed and predicted values always between 3% and 10%. Predictions are slightly better under drought condition because this condition is the reference condition in the parameterization of the model, so the effects linked to this condition are estimated with more observations than the normal condition. Predictions are also less reliable for the smallest and the largest heat times. The residual variance is homoscedastic in the chosen model so that at the start of growth (*i*.*e*. small heat times) where the height measurements are small, the noise is proportionally greater, explaining the slightly poorer predictions for this part of the growth curve. On the other hand, the model predicts a plateau at the end of the growth curve, from which certain observations deviate over the last heat time, as shown in Figure 11. This may explain the slight deterioration in predictions at the end of the curve. Improving the model to better fit the start and end of the growth curve would certainly further improve the quality of predictions.

**Fig 13.**
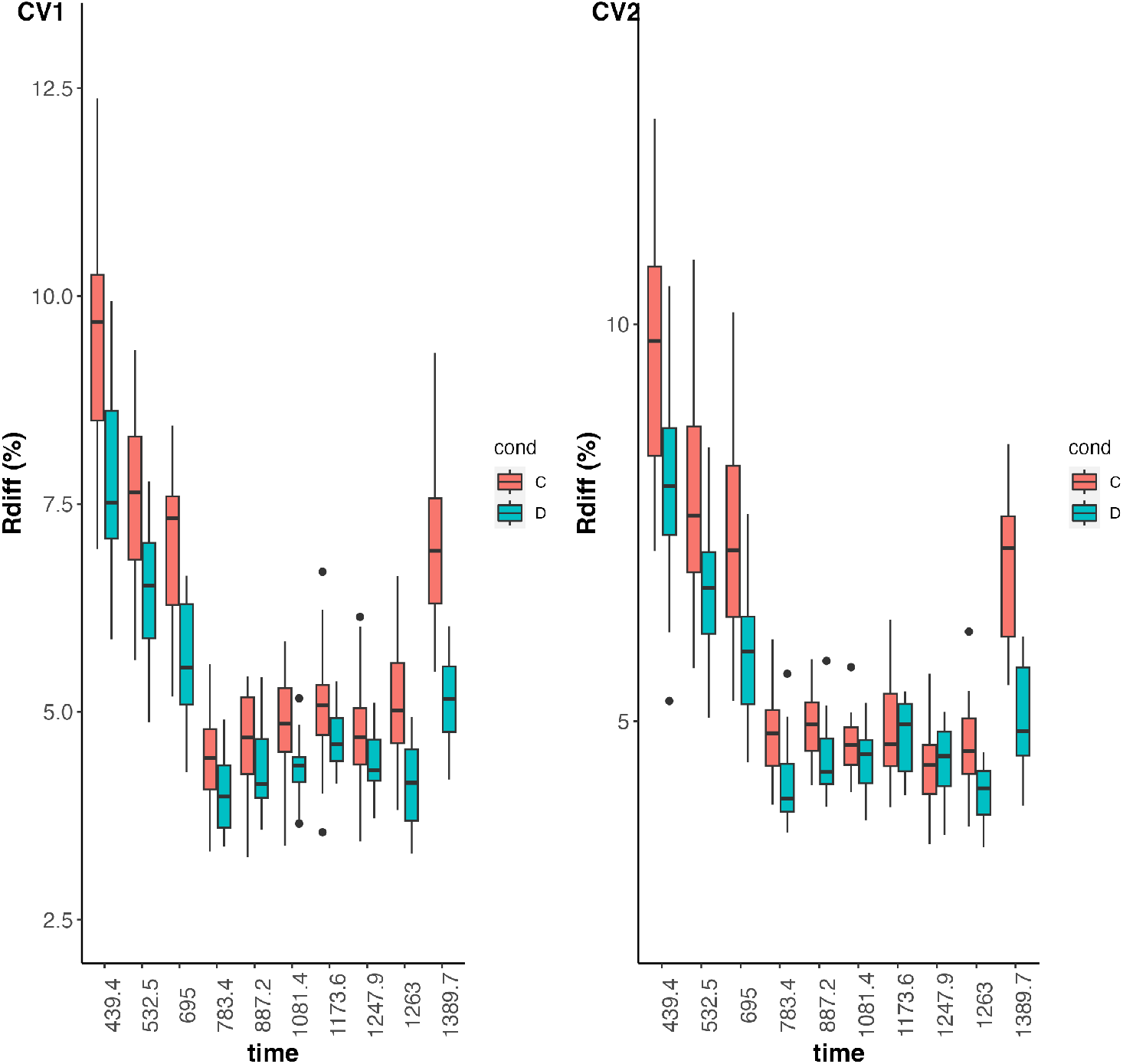
Boxplots of relative differences (%) between predicted and observed height measures in the validation samples per heat time point and hydric condition (C: normal hydric condition, D: drought condition) computed by cross-validation under cv1 (left) and cv2 (right). Smaller relative difference is better.

Note that in the present study, only 4 plants were observed for each variety. As suggested by the simulation study, the overall robustness of the analysis could be improved by increasing the number of plants per variety to provide better information on the differences between the varieties.

#### 3.2.2 Comparison with the Genome Based Model method

We compared our approach to the Genome Based Model method described in [15]. To do this, we used the `GenomeBasedModel` package in `R`, available at the following link https://github.com/Onogi/GenomeBasedModel. We have limited the comparison to predictive performance under the cv1 scenario, where we seek to make predictions on varieties that have not been observed in any environment, reproducing the cross-validation study on the same training and validation samples extracted from real data as above. The results in terms of relative differences between predicted and observed values are shown in Figure 14 and show that comparable performance is achieved with both approaches. There are, however, fundamental differences in terms of modeling between the two approaches, that make our approach much more informative. In particular, the Genome Based Model method does not take into account a genetic covariance matrix, *G* in the present work, and therefore forces us to consider independent nonlinear model parameters for each unit.

**Fig 14.**
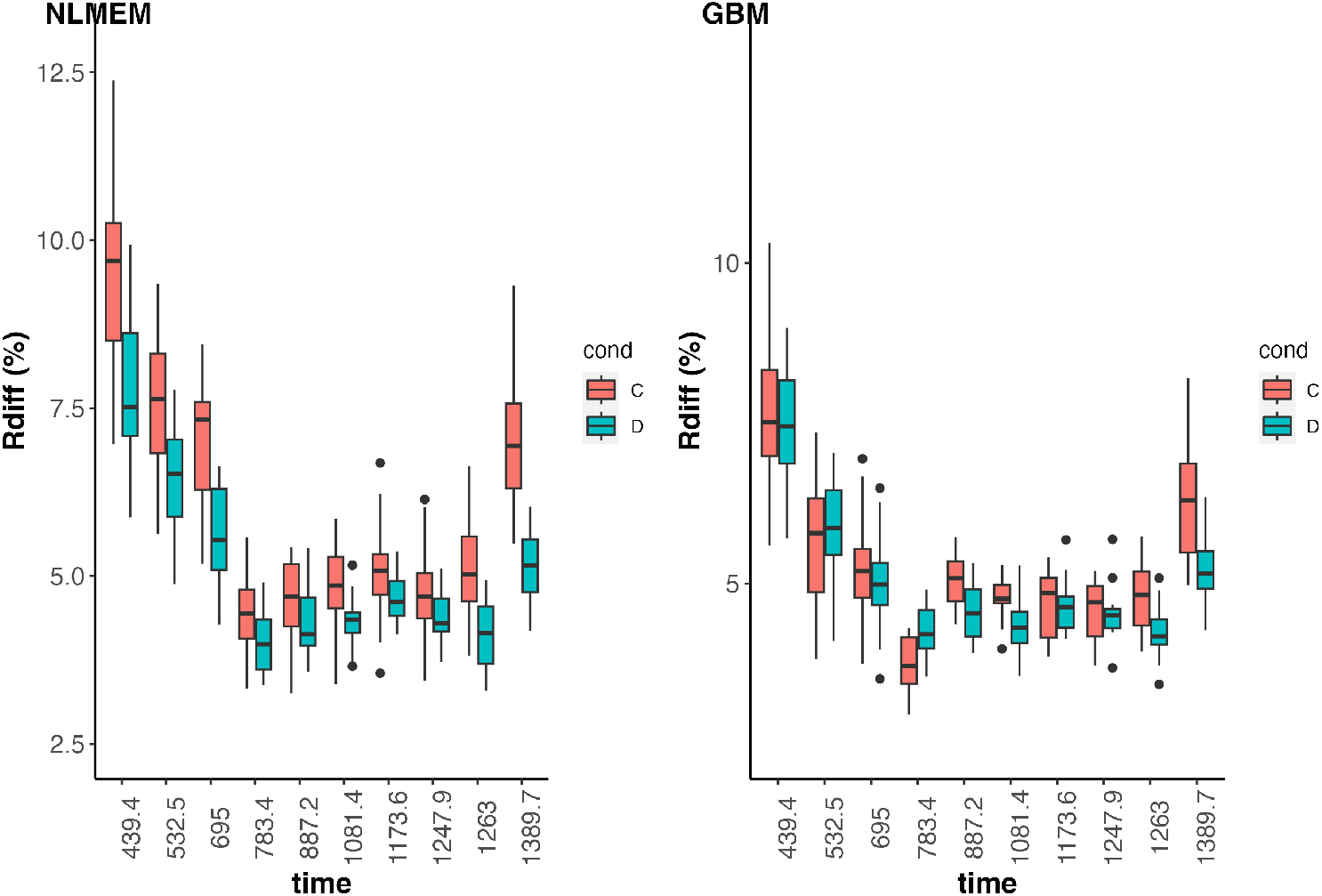
Boxplots of relative differences (%) between predicted and observed height measures in the validation samples per heat time point and hydric condition (C: normal hydric condition, D: drought condition) computed by cross-validation under cv1. Left panel depicts the results obtained with the NLMEM approach and right panel depicts the results obtained with the GBM method [15]. Smaller relative difference is better.

## 4 Discussion

Nonlinear models have been used in the genomic prediction modeling of plant and animal growth [6, 12, 13, 33, 34]. In genome prediction modeling, a fusion of nonlinear functions and mixed-effects models, that is, the NLMEM framework, is required. Two approaches have been used: a two-step approach in which a nonlinear function is fitted beforehand and its parameters are modeled by a mixed-effects model [12, 13], and a one-step approach in which these are performed in a single step [6, 33, 34]. The two-step approach requires estimating the parameters of the nonlinear function for each plant, sometimes resulting in overfitting. The one-step approach, in contrast, has the advantage of robust estimation because the parameters are estimated simultaneously for all plants considering genetic relationships and differences in cultivation treatments. In this study, we applied the NLMEM framework using a one-step approach and modeled the growth of numerous soybean varieties under different environmental conditions using a single model. The NLMEM framework allowed us to separate the variation in the growth model parameters into genetic and environmental variations as well as the effect of growing conditions (irrigation or no irrigation) on these variations.

Whole-genome DNA polymorphisms of soybean varieties have been decoded and classified into three groups based on the genetic similarities between varieties [41]. The genetic variation in the parameters estimated in this study clearly showed that there were differences in growth patterns among these classified groups. The “Primitive” varieties, which were considered close to the soybean varieties when they were first introduced to cultivation, showed fast and vigorous growth, while the varieties belonging to the “Japan” group exhibited the opposite patterns. However, these patterns are average trends, and some varieties in the same group showed different patterns. For example, some of the “Japan” varieties showed fast and vigorous growth. The growth pattern of soybeans is also related to weed suppression [48], which is an important trait in breeding. Characterization of genetic variations in growth patterns using the method proposed in this study is expected to be effective in soybean breeding.

The proposed method has the following characteristics compared to previous one-step approaches [6, 33, 34]. First, the parameters of the nonlinear model, which can be regarded as latent variables, can be calculated with high speed and efficiency using an efficient algorithm called the SAEM, which is an extension of the EM algorithm. In contrast, previous methods based on the one-step approach used Markov chain Monte Carlo (MCMC) and variational Bayesian methods. The proposed inference methodology with SAEM has several advantages over competing methods. First, the SAEM algorithm is less computationally intensive and much faster than the MCMC implementations. A thorough comparison between SAEM and MCMC was performed by [44] for Bayesian variable selection in NLMEMs, which highlighted the faster execution times associated with SAEM than with MCMC. In practice, the computation times vary with sample characteristics and model complexity. For the soybean data example processed with R software with a 2GHz intel core i5 processor, parameter estimation lasted 17 min and genetic effects prediction lasted 47 min, which is very promising compared with the computation times reported in the literature for similar analyses.

Another feature of the proposed method is that it considers not only the genetic correlation among parameters but also the correlation structure in the error. In the previous one-step approach-based methods, estimation was performed assuming that the errors were independent of the parameters [6, 33]. In this study, by assuming non-independence of errors among parameters. Based on the estimated genetic and environmental variance-covariance matrices of the growth model parameters, we were able to determine not only the heritability of each parameter but also the genetic and phenotypic correlations (correlations due to both genetic and environmental effects) among the parameters. Such information can be used to determine the degree of genetic control of growth and predict selection responses when selection is based on growth patterns. Genetic correlation can also be an indicator of the difficulty of simultaneously modifying two parameters in a certain direction. By comparing phenotypic and genetic correlations, it is possible to ascertain the extent to which the correlation that appears as phenotypic is due to inheritance.

In addition to the logistic growth and asymptotic regression curves used in this study, a variety of nonlinear models are available [12, 49]. Various covariance matrices representing the genetic similarity among varieties have also been used in addition to the additive genomic relationship matrix used in this study, including a pedigree-based relationship matrix and a nonlinear relationship matrix, such as a dominance relationship matrix and a Gaussian kernel [32]. Further, in this study, we used two levels of irrigation as part of the cultivation management treatment, but various treatments and levels are possible [50]. The proposed methodology can be straightforwardly adapted to different contexts without any major impact on the general implementation of the SAEM algorithm, by changing either the nonlinear model function *g*(·) to better reflect the curve profiles or the definition of matrices *X*_*i*_ and *Z*_*i*_ depending on the number and the potential fixed and genetic effects of the available covariates. In particular, it is very easy to consider more than two environmental conditions using the proposed approach. More generally, our method can be applied to any nonlinear model *g*(*φ, t*), whether it is an empirical, mechanistic or ecophysiological model, as long as *g*(*φ, t*) can be numerically evaluated at any value of *φ* and *t*. Of course, the execution time of the algorithms we propose in the present paper depends on the time required to evaluate the model *g*, which is used at each iteration. However, it should be highlighted that, as shown in our data analysis, a careful choice of *g*(·) is crucial to ensure good estimates and predictions. Users should also note that the performance of the modeling approach based on NLMEM depends on the structure of the collected data, as demonstrated in the simulation study.

As shown in the simulated data study, by varying the number of varieties and plants and the data collection scheme, observing enough varieties, plants per variety, and points on the growth curves is essential to ensure accurate parameter estimations and predictions of the genetic effects. While one can reasonably trust the results obtained on the soybean growth data analyzed in this paper, as suggested by the simulated results on datasets with similar structures, the data collection must be streamlined beforehand to be able to collect data of sufficient quality and ensure that the NLMEM approach provides reliable and robust results. Note that during the simulations, data from 2000 plants were used simultaneously to estimate parameters. This also demonstrates the scalability of the proposed method for large data-sets.

Recently, remote sensing has enabled the highly efficient measurement of plant and animal growth over time. Furthermore, the determination of genome-wide markers has become more efficient, making it possible to measure both genome-wide polymorphisms and growth for a large number of genotypes. Combining the measurements obtained from such technological advances with the methods proposed in this study could genetically model the growth patterns of plants and animals, and help to understand, control, and predict their inheritance. The method proposed in this study is scalable to larger data sets owing to its computational speed. The codes for the method and the data used in this study are available at https://github.com/madelattre/Scripts-soybean-paper and can be extended according to the conditions of the application in each research project.

## Supporting information

### Computer codes

The codes are available at https://github.com/madelattre/Scripts-soybean-paper

## S1 Appendix.

**Algorithmic details**

The objective of this appendix is to provide the main technical elements for implementing the SAEM algorithm in model (1)-(2). First, we introduce the compact matrix notations:

- 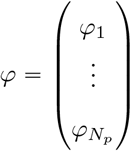: the *m*.*N*_*p*_ vector obtained by concatenation of the *φ*_*i*_’s,
- 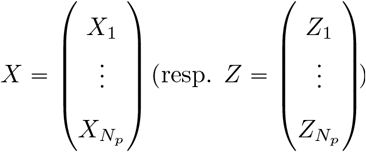: the *m*.*N*_*p*_ × *p* (resp. *m*.*N*_*p*_ × *q*.*N*_*v*_) matrix obtained by vertical concatenation of the *X*_*i*_’s (resp. *Z*_*i*_’s),

1. **Key distributions in model** (1)**-**(2)
  a. **Complete data log-likelihood**. Using that in model (1)-(2), 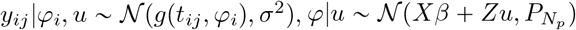 and *u* ∼ (**0**, *K* ⨂ *G*), we get

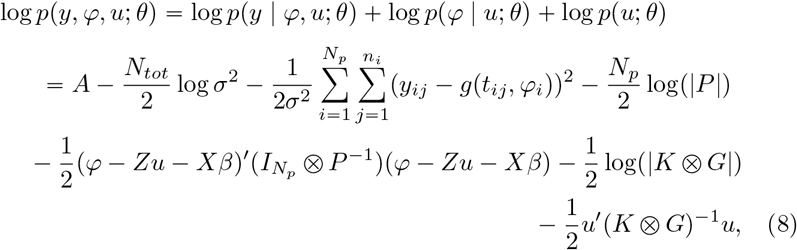

where *A* is a constant term which does not depend on *θ*, 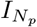 is the identity matrix of size *N*_*p*_, *n*_*i*_ is the number of observations for plant *i* ∈ {1, …, *N*_*p*_} and 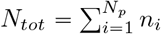 is the total number of observations available in the sample (*i*.*e*. all observations of all plants).
  b. **Conditional distribution of the genetic effects** *u* **given** *φ*. Using the Bayes formula where *p*(*u*|*φ θ*) = *p*(*φ*|*u*; *θ*)*p*(*u*; *θ*){ (*φ*; *θ*) and that in model (1)-(2), 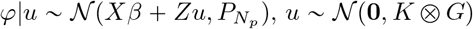 and 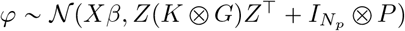, by elementary calculations, we get that *u*|*φ* ∼ 𝒩 (*m*_*u*_(*θ*), Σ_*u*_(*θ*)), where

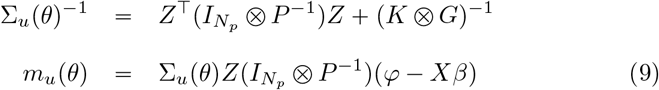
2. **SAEM algorithm** Algorithm 1 describes the algorithm used to estimate parameters. *K* is the total number of iterations, which is tuned by the user such that the algorithm converges at the end of *K* iterations. The stochastic approximation step uses model (1)-(2) belonging to the curved exponential family, and thus consists of the computation of the stochastic approximations of the sufficient statistics (see [39]). **Remarks:**
  - To better manage the acceptance proportions at the simulation step of the algorithm, it is possible to perform several (but a finite number of) iterations of Metropolis-Hastings rather than one, and define 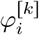 as the last accepted value after these iterations.
  - Choosing *q*_*i,θ*_(*φ*) as *p*(*φ*|*u*; *θ*) allows to simplify the expression of the acceptance ratio *α*_*i,k*_.
  - In some situations, in particular when the number of observed plants per variety is small, the convergence of the algorithm can be improved by using *M* Monte-Carlo Markov chains instead of only one as described in Algorithm 1. This amounts to simulating *M* values 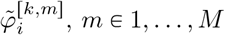, at each iteration and replacing the sufficient statistics by their means over the *M* simulated values.

Algorithm 2 presents algorithm for genetic effects prediction. This should be used with 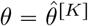 obtained at the convergence of the first algorithm. In general, Algorithm 2 is written for any value of *θ*. It also exploits the fact that the model (1)-(2) belongs to the curved exponential family.

## S2 Appendix.

**Convergence graphs**.

### Algorithm 1

SAEM algorithm for parameter estimation

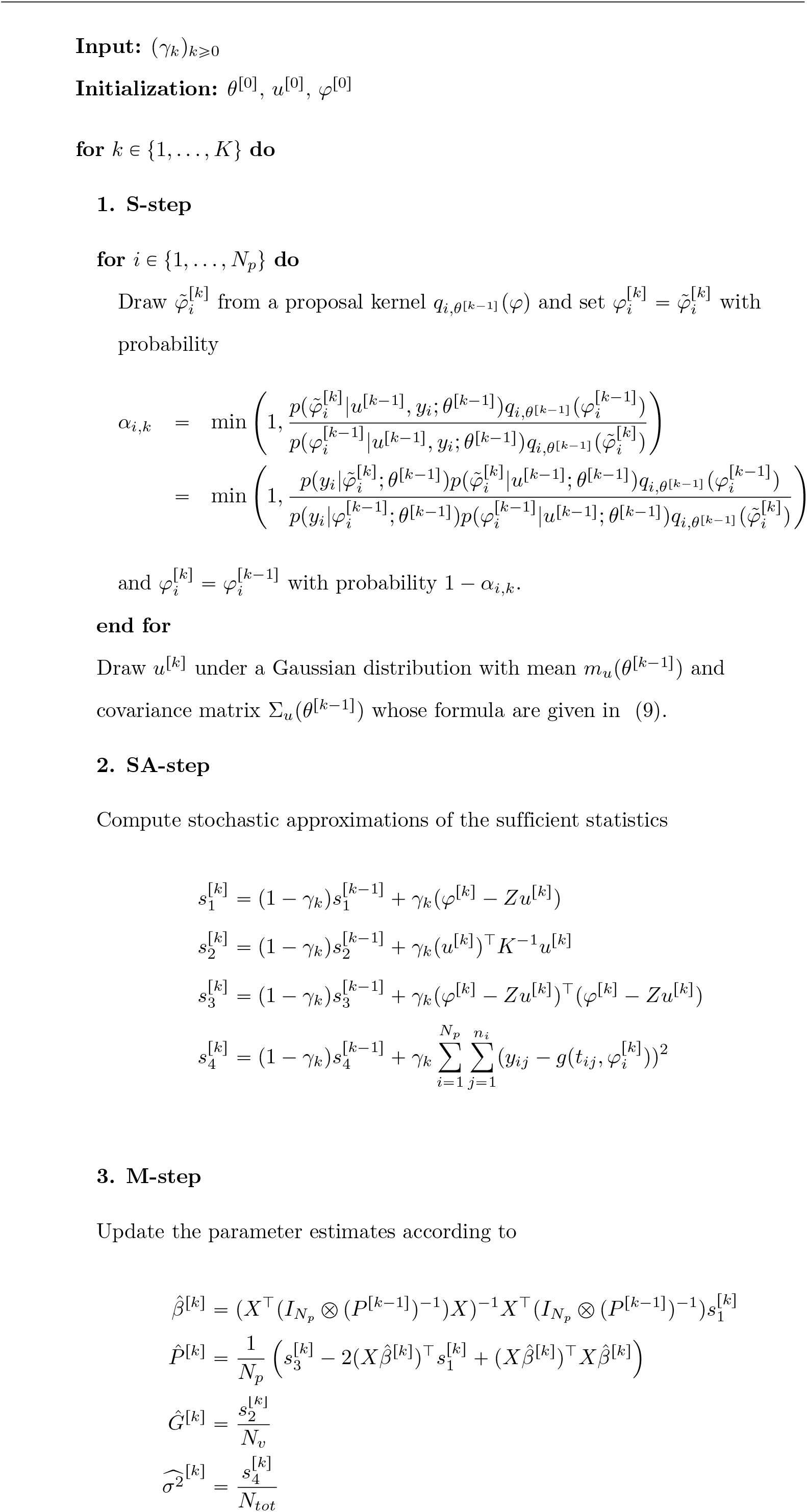

### Algorithm 2

SAEM algorithm for genetic effects prediction

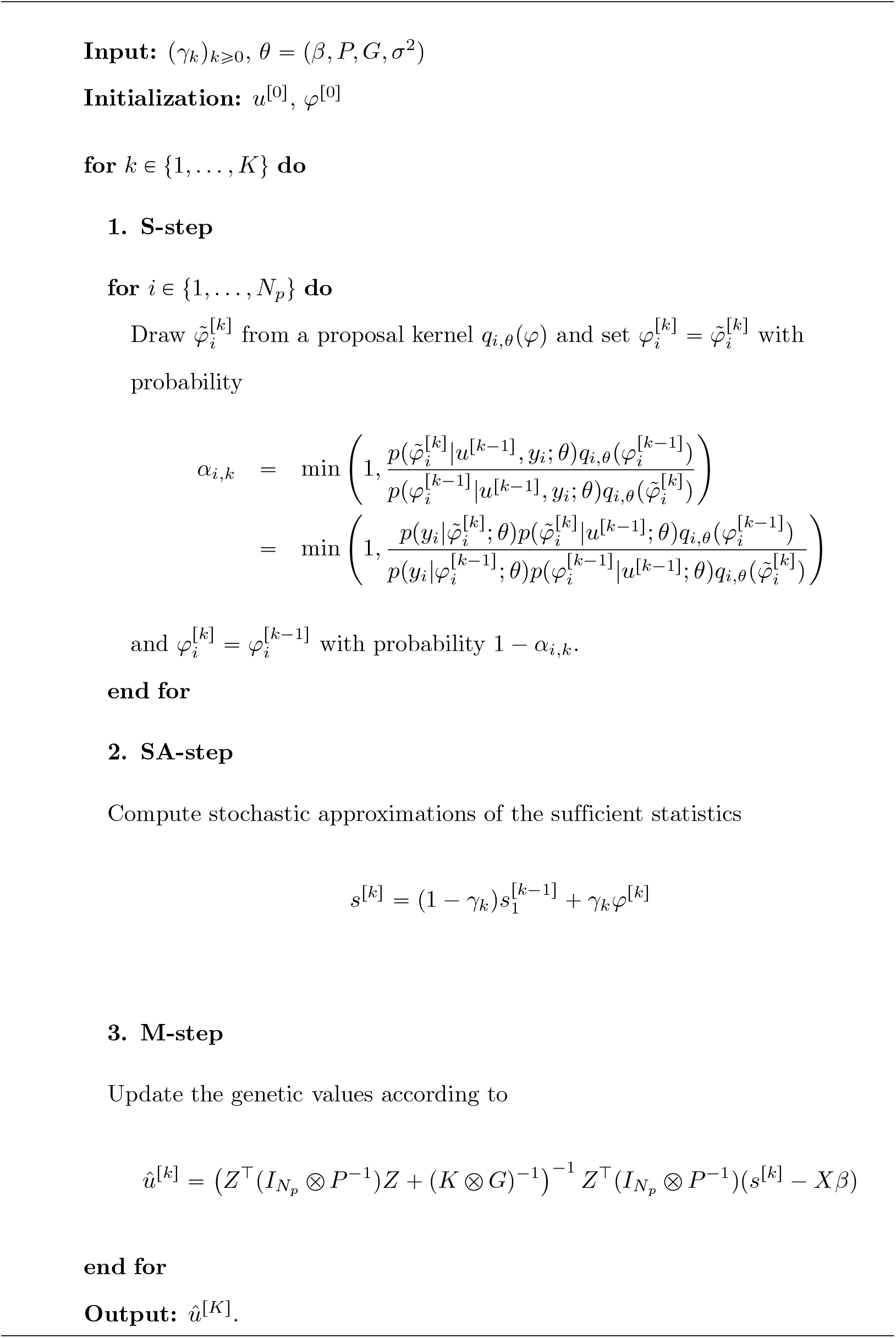

## Acknowledgments

This study was supported by JST CREST (grant no. JPMJCR16O2) and JSPS KAKENHI (grant no. JP22H02306). The funders had no role in the study design, data collection and analysis, decision to publish, or manuscript preparation.

We are grateful to the technical staff of the Arid Land Research Center, Tottori University, and Izumi Higashida for the management of the field experiments. We would like to thank Yoshihiro Ohmori, Yuji Yamasaki, Hirokazu Takahashi, Hideki Takanashi, Mai Tsuda, Yuji Sawada, Hisashi Tsujimoto, Akito Kaga, Mikio Nakazono, Toru Fujiwara for being involved in the field experiments, Hiromi Kajiya-Kanegae and Kosuke Hamazaki for curating whole-genome marker data, and Sawako Maruyama for curating the phenotype data.

## List of Figures

**Fig 15.**
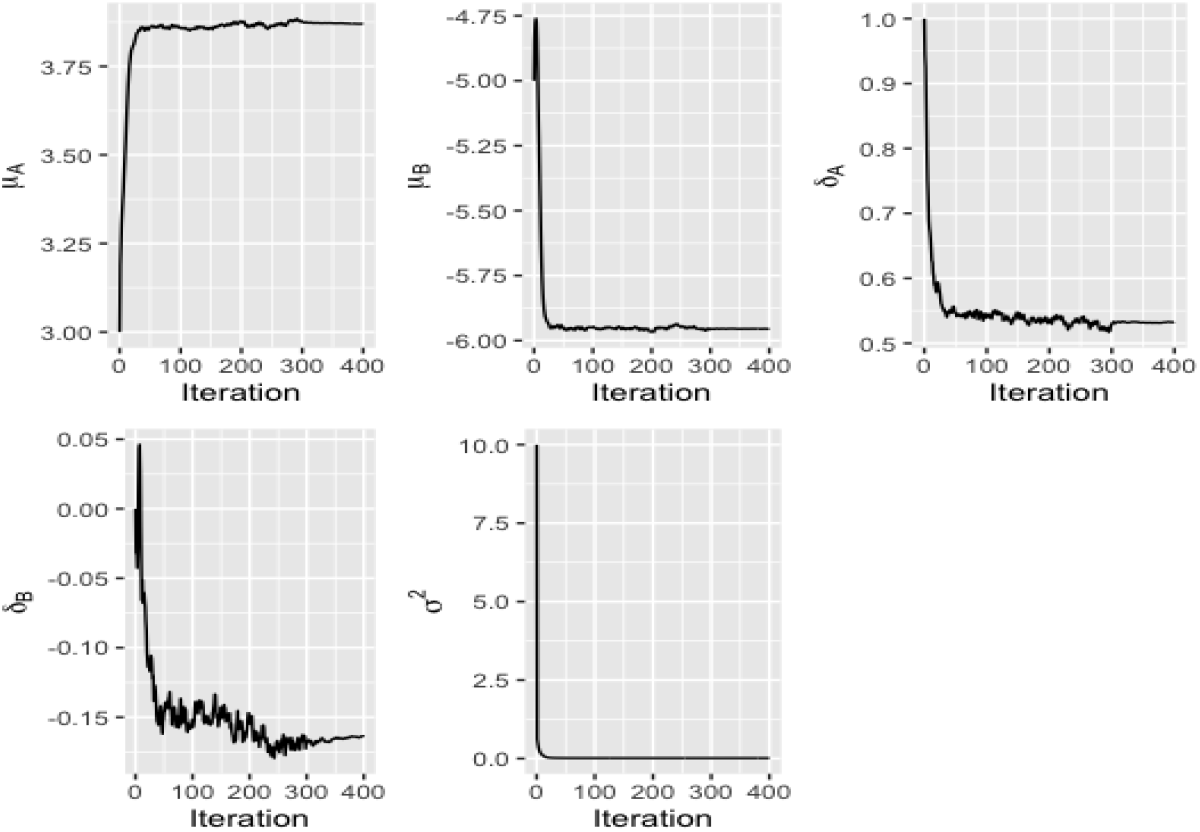
Real data analysis: Parameter estimation. Evolution of *β*’s and *σ*^2^’s estimations over iterations of the SAEM algorithm.

**Fig 16.**
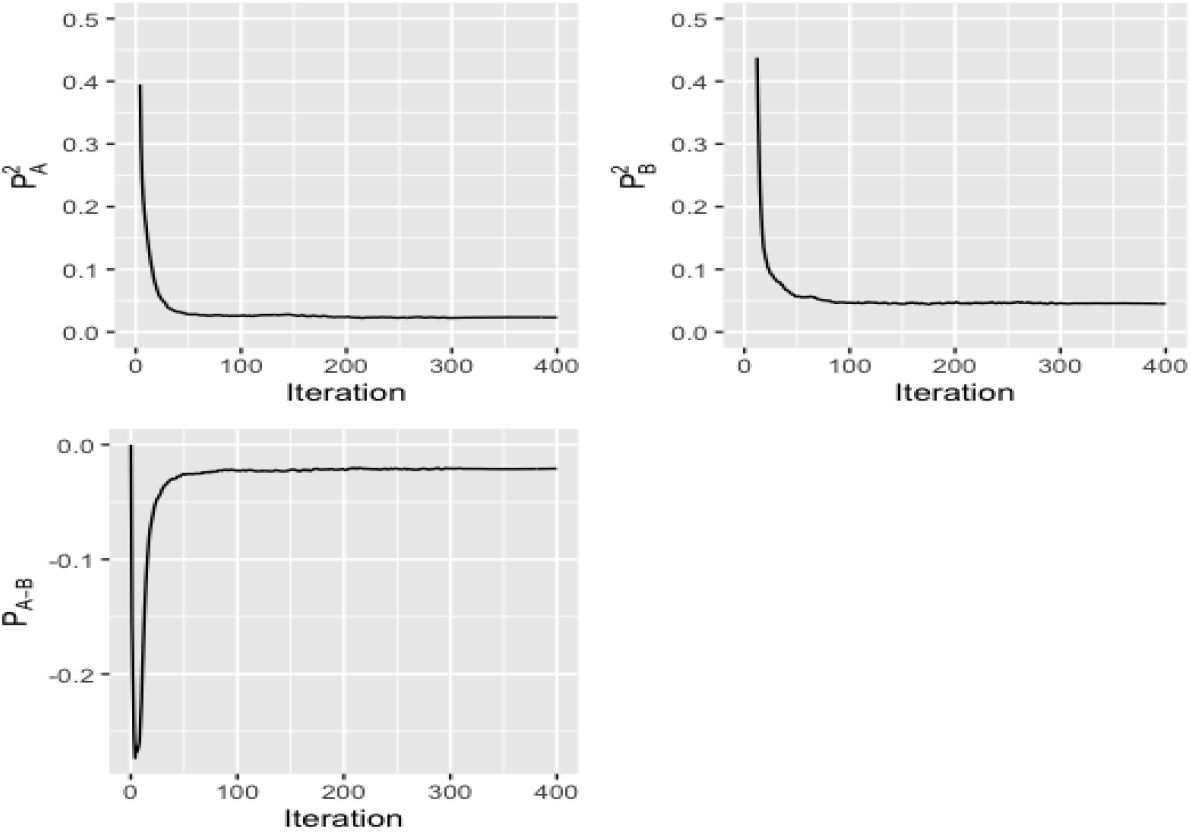
Real data analysis: Parameter estimation. Evolution of *P* ‘s estimation over iterations of the SAEM algorithm.

**Fig 17.**
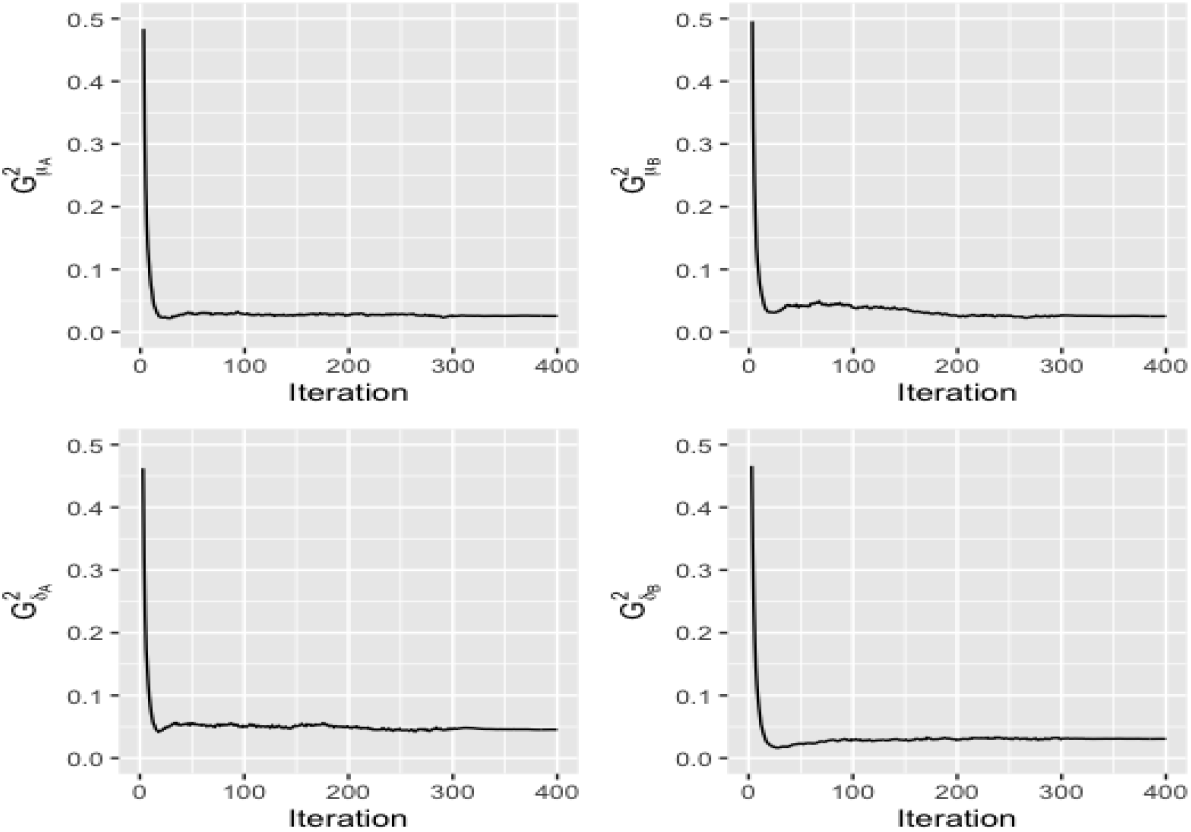
Real data analysis: Parameter estimation. Evolution of the estimation of the diagonal terms of *G* over iterations of the SAEM algorithm.

**Fig 18.**
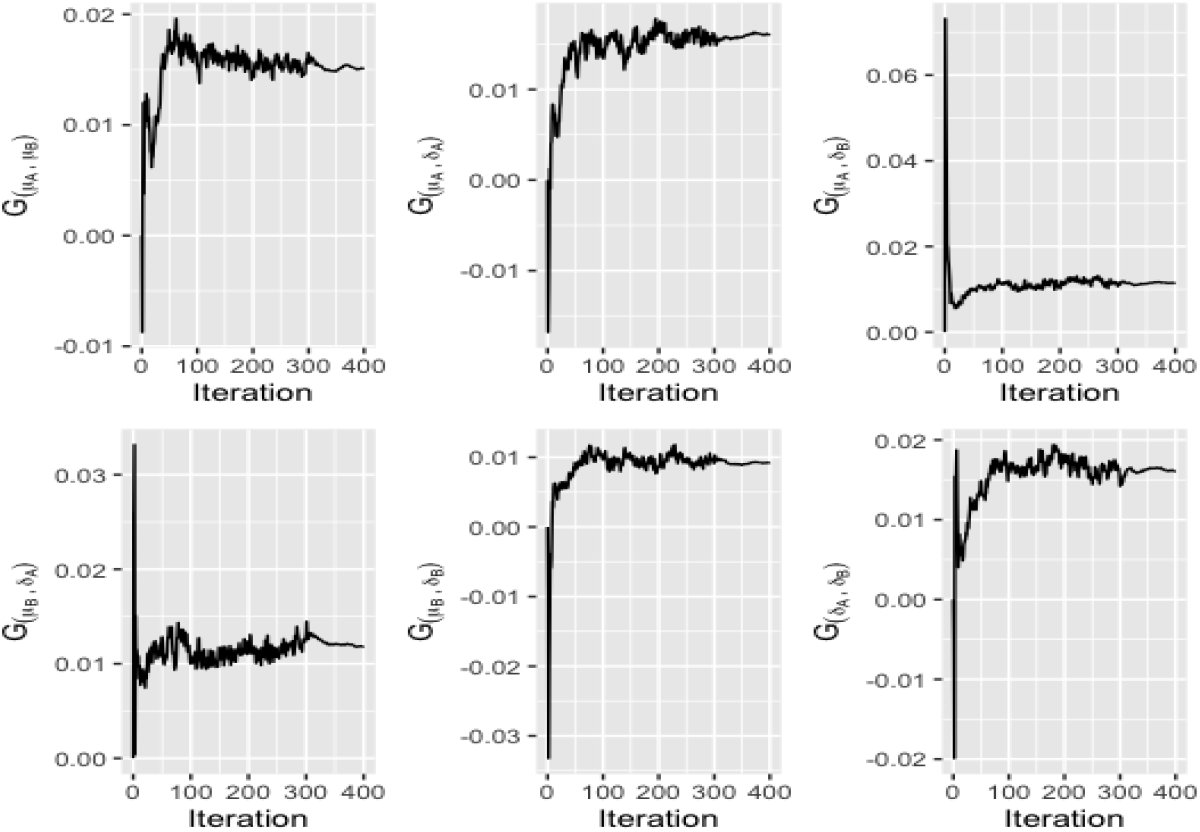
Real data analysis: Parameter estimation. Evolution of the estimation of the extra-diagonal terms of *G* over iterations of the SAEM algorithm.

## References

1. Soltani A. Modeling physiology of crop development, growth and yield. CABi; 2012.

2. Yang G, Liu J, Zhao C, Li Z, Huang Y, Yu H, et al. Unmanned Aerial Vehicle Remote Sensing for Field-Based Crop Phenotyping: Current Status and Perspectives. Frontiers in Plant Science. 2017;8:1111. doi:10.3389/fpls.2017.01111.

3. White JW, Andrade-Sanchez P, Gore MA, Bronson KF, Coffelt TA, Conley MM, et al. Field-based phenomics for plant genetics research. Field Crops Research. 2012;133:101–112. doi:10.1016/j.fcr.2012.04.003.

4. Andrade-Sanchez P, Gore MA, Heun JT, Thorp KR, Carmo-Silva AE, French AN, et al. Development and evaluation of a field-based high-throughput phenotyping platform. Functional Plant Biology. 2013;41(1):68–79. doi:10.1071/fp13126.

5. Campbell M, Walia H, Morota G. Utilizing random regression models for genomic prediction of a longitudinal trait derived from highthroughput phenotyping. Plant Direct. 2018;2(9):e00080. doi:10.1002/pld3.80.

6. Campbell MT, Grondin A, Walia H, Morota G. Leveraging genome-enabled growth models to study shoot growth responses to water deficit in rice. Journal of experimental botany. 2020;71(18):5669–5679.

7. Toda Y, Kaga A, KajiyaKanegae H, Hattori T, Yamaoka S, Okamoto M, et al. Genomic prediction modeling of soybean biomass using UAVbased remote sensing and longitudinal model parameters. The Plant Genome. 2021; p. e20157. doi:10.1002/tpg2.20157.

8. Nelder J. The fitting of a generalization of the logistic curve. Biometrics. 1961;17(1):89–110.

9. Winsor CP. The Gompertz curve as a growth curve. Proceedings of the national academy of sciences. 1932;18(1):1–8.

10. Crispim AC, Kelly MJ, Guimaraes SEF, Silva FFe, Fortes MRS, Wenceslau RR, et al. Multi-Trait GWAS and New Candidate Genes Annotation for Growth Curve Parameters in Brahman Cattle. PLOS ONE. 2015;10(10):e0139906. doi:10.1371/journal.pone.0139906.

11. Baker R, Leong W, Welch S, Weinig C. Mapping and predicting non-linear Brassica rapa growth phenotypes based on Bayesian and frequentist complex trait estimation. G3: Genes, Genomes, Genetics. 2018;8(4):1247–1258.

12. Yin T, Konig S. Genomic predictions of growth curves in Holstein dairy cattle based on parameter estimates from nonlinear models combined with different kernel functions. Journal of dairy science. 2020;103(8):7222–7237.

13. Toda Y, Sasaki G, Ohmori Y, Yamasaki Y, Takahashi H, Takanashi H, et al. Genomic prediction of green fraction dynamics in soybean using unmanned aerial vehicles observations. Frontiers in Plant Science. 2022;13.

14. Meuwissen TH, Hayes BJ, Goddard ME. Prediction of total genetic value using genome-wide dense marker maps. Genetics. 2001;157(4):1819–29.

15. Onogi A. Connecting mathematical models to genomes: Joint estimation of model parameters and genome-wide marker effects on these parameters. Bioinformatics. 2020;36(10):3169–3176. doi:10.1093/bioinformatics/btaa129.

16. Henderson Jr CR. Analysis of covariance in the mixed model: higher-level, nonhomogeneous, and random regressions. Biometrics. 1982; p. 623–640.

17. Laird NM, Ware JH. Random-effects models for longitudinal data. Biometrics. 1982; p. 963–974.

18. Schaeffer LR. Application of random regression models in animal breeding. Livestock Production Science. 2004;86(1):35–45. doi:10.1016/S0301-6226(03)00151-9.

19. Oliveira H, Brito L, Lourenco D, Silva F, Jamrozik J, Schaeffer L, et al. Invited review: Advances and applications of random regression models: From quantitative genetics to genomics. Journal of dairy science. 2019;102(9):7664–7683.

20. Campbell M, Walia H, Morota G. Utilizing random regression models for genomic prediction of a longitudinal trait derived from high-throughput phenotyping. Plant Direct. 2018;2(9):e00080.

21. Sun J, Rutkoski JE, Poland JA, Crossa J, Jannink JL, Sorrells ME. Multitrait, random regression, or simple repeatability model in high-throughput phenotyping data improve genomic prediction for wheat grain yield. The plant genome. 2017;10(2):plantgenome2016–11.

22. Technow F, Messina CD, Totir LR, Cooper M. Integrating crop growth models with whole genome prediction through approximate Bayesian computation. PloS one. 2015;10(6):e0130855.

23. Onogi A, Watanabe M, Mochizuki T, Hayashi T, Nakagawa H, Hasegawa T, et al. Toward integration of genomic selection with crop modelling: the development of an integrated approach to predicting rice heading dates. Theoretical and Applied Genetics. 2016;129(4):805–817. doi:10.1007/s00122-016-2667-5.

24. Cooper M, Technow F, Messina C, Gho C, Totir LR. Use of Crop Growth Models with Whole-Genome Prediction: Application to a Maize Multienvironment Trial. Crop Science. 2016;56(5):2141–2156. doi:10.2135/cropsci2015.08.0512.

25. Messina CD, Technow F, Tang T, Totir R, Gho C, Cooper M. Leveraging biological insight and environmental variation to improve phenotypic prediction: Integrating crop growth models (CGM) with whole genome prediction (WGP). European Journal of Agronomy. 2018;100:151–162. doi:10.1016/j.eja.2018.01.007.

26. Diepenbrock CH, Tang T, Jines M, Technow F, Lira S, Podlich D, et al. Can we harness digital technologies and physiology to hasten genetic gain in US maize breeding? Plant Physiology. 2021;188(2):1141–1157. doi:10.1093/plphys/kiab527.

27. Onogi A. In: Ahmadi N, Bartholome J, editors. Integration of Crop Growth Models and Genomic PredictionGenomic predictions (GP). New York, NY: Springer US; 2022. p. 359–396. Available from: 10.1007/978-1-0716-2205-6_13.

28. Poudel P, Naidenov B, Chen C, Alderman PD, Welch SM. Integrating genomic prediction and genotype specific parameter estimation in ecophysiological models: overview and perspectives. in silico Plants. 2023;5(1):diad007. doi:10.1093/insilicoplants/diad007.

29. Ma CX, Casella G, Wu R. Functional Mapping of Quantitative Trait Loci Underlying the Character Process: A Theoretical Framework. Genetics. 2002;161(4):1751–1762. doi:10.1093/genetics/161.4.1751.

30. Wu R, Ma CX, Lin M, Casella G. A General Framework for Analyzing the Genetic Architecture of Developmental Characteristics. Genetics. 2004;166(3):1541–1551. doi:10.1534/genetics.166.3.1541.

31. Gianola D. Priors in Whole-Genome Regression: The Bayesian Alphabet Returns. Genetics. 2013;194(3):573–596. doi:10.1534/genetics.113.151753.

32. Morota G, Gianola D. Kernel-based whole-genome prediction of complex traits: a review. Frontiers in Genetics. 2014;5:363. doi:10.3389/fgene.2014.00363.

33. Onogi A, Ogino A, Sato A, Kurogi K, Yasumori T, Togashi K. Development of a structural growth curve model that considers the causal effect of initial phenotypes. Genetics Selection Evolution. 2019;51(1):19.

34. Yu H, van Milgen J, Knol E, Fernando R, Dekkers J. A Bayesian hierarchical model to integrate a mechanistic growth model in genomic prediction. In: 12. World congress on genetics applied to livestock production (WCGALP); 2022.

35. Pinheiro J C, Bates D M. Mixed-Effects Models in S and S-PLUS. 1st ed. Statistics and Computing. Springer-Verlag New York Inc.; 2000.

36. Lavielle M. Mixed Effects Models for the Population Approach. Models, Tasks, Methods and Tools. Chapman and Hall/CRC Biostatistics Series. Chapman and Hall/CRC; 2014.

37. Jaffrezic F, Meza C, Lavielle M, Foulley JL. Genetic analysis of growth curves using the SAEM algorithm. Genetics Selection Evolution. 2006;38(6):583–600.

38. Dempster A P, Laird N M, Rubin D B. Maximum Likelihood from Incomplete Data via the EM Algorithm. Journal of Royal Statistical Society Series B. 1977;39(1):1–38.

39. Delyon B, Lavielle M, Moulines E. Convergence of a stochastic approximation version of the EM algorithm. The Annals of Statistics. 1999;27(1):94–128.

40. Kaga A, Shimizu T, Watanabe S, Tsubokura Y, Katayose Y, Harada K, et al. Evaluation of soybean germplasm conserved in NIAS genebank and development of mini core collections. Breeding science. 2012;61(5):566–592.

41. Kajiya-Kanegae H, Nagasaki H, Kaga A, Hirano K, Ogiso-Tanaka E, Matsuoka M, et al. Whole-genome sequence diversity and association analysis of 198 soybean accessions in mini-core collections. DNA Research. 2021;28(1).

42. Browning BL, Zhou Y, Browning SR. A One-Penny Imputed Genome from Next-Generation Reference Panels. The American Journal of Human Genetics. 2018;103(3):338–348. doi:10.1016/j.ajhg.2018.07.015.

43. Stekhoven DJ. missForest: Nonparametric Missing Value Imputation using Random Forest; 2022. https://cran.r-project.org/web/packages/missForest/index.html.

44. Naveau M, King GKK, Rincent R, Sansonnet L, Delattre M. Bayesian high-dimensional covariate selection in non-linear mixed-effects models using the SAEM algorithm; 2022.

45. Kuhn E, Lavielle M. Coupling a stochastic approximation version of EM with an MCMC procedure. ESAIM:PS. 2004;8:115–131.

46. Robert CP, Casella G, Casella G. Monte Carlo statistical methods. vol. 2. Springer; 1999.

47. Costa-Neto G, Fritsche-Neto R, Crossa J. Nonlinear kernels, dominance, and envirotyping data increase the accuracy of genome-based prediction in multi-environment trials. Heredity. 2021;126(1):92–106.

48. Kurokawa S, Kaga A, Tsuda M, Sekine D, Shibuya T. Evaluation of validity and limitations of the soybean canopy height-to-row spacing ratio as an onsite index to control weeds using diverse soybean accessions. Weed Biology and Management. 2019;19(3):103–110.

49. Hossein-Zadeh NG. Modelling growth curve in Moghani sheep: comparison of non-linear mixed growth models and estimation of genetic relationship between growth curve parameters. The Journal of Agricultural Science. 2017;155(7):1150–1159.

50. Sakurai K, Toda Y, Kajiya-Kanegae H, Ohmori Y, Yamasaki Y, Takahashi H, et al. Time-series multispectral imaging in soybean for improving biomass and genomic prediction accuracy. The Plant Genome. 2022;15(4):e20244.

